# Estimating the burden of α-thalassaemia in Thailand using a comprehensive prevalence database for Southeast Asia

**DOI:** 10.1101/412718

**Authors:** Carinna Hockham, Supachai Ekwattanakit, Samir Bhatt, Bridget S Penman, Sunetra Gupta, Vip Viprakasit, Frédéric B Piel

## Abstract

Severe forms of α-thalassaemia, haemoglobin H disease and haemoglobin Bart’s hydrops fetalis, are an important public health concern in Southeast Asia. Yet information on the prevalence, genetic diversity and health burden of α-thalassaemia in the region remains limited. We compiled a geodatabase of α-thalassaemia prevalence and genetic diversity surveys and, using geostatistical modelling methods, generated the first continuous maps of α-thalassaemia mutations in Thailand and sub-national estimates of the number of newborns with severe forms in 2020. We also summarised the current evidence-base for α-thalassaemia prevalence and diversity for the region. We estimate that 3,595 (95% credible interval 1,717 – 6,199) newborns will be born with severe α-thalassaemia in Thailand in 2020, which is considerably higher than previous estimates. Accurate, fine-scale epidemiological data are necessary to guide sustainable national and regional health policies for α-thalassaemia control. Our maps and newborn estimates are an important first step towards this aim.

**Funding:** This work was supported by European Union’s Seventh Framework Programme (FP7//2007-2013)/European Research Council [268904 – DIVERSITY]

## Introduction

α-thalassaemia is one of the commonest monogenic disorders of humans, spanning much of the malaria belt, including the Mediterranean, sub-Saharan Africa, Asia and the Pacific. It is estimated that up to 5% of the world’s population carries at least one α-thalassaemia variant, with some populations in Southeast Asia reporting gene frequencies of close to 80% (1). Central to its elevated frequency is the malaria protection afforded by the underlying genetic mutations, which have been favoured by natural selection in populations with historically high rates of malaria (2-5). Due to recent population migrations, α-thalassaemia is now common in other parts of the world, as illustrated by the inclusion of haemoglobin H (HbH) disease (a form of α-thalassaemia) in the newborn screening programme in California (6, 7).

Humans typically possess four copies of the α-globin gene. In an individual with α-thalassaemia, at least one of these four copies is absent or dysfunctional. The resulting deficit in α-globin affects the balance between α-globin and β- or γ-globin chains that is necessary to produce normal adult haemoglobin (HbA) and normal foetal haemoglobin (HbF), respectively (8). The severity of α-thalassaemia is inversely related to the number of functional copies of the α-globin gene. A deficit of three or more α-globin genes leads to the production of γ-globin tetramers, called Hb Bart’s, in the foetus or β-globin tetramers, called HbH, in adults. Due to their very high oxygen affinity, neither tetramer is capable of transporting oxygen efficiently (9). Furthermore, the instability of HbH leads to the production of inclusion bodies in red blood cells and a variable degree of haemolytic anaemia.

To date, 121 α-globin gene mutations have been identified (HbVar, http://globin.bx.psu.edu, accessed 07 July 2018). These include: (i) double gene deletions that remove both α-globin copies in a gene pair (α^0^-thalassaemia), (ii) single gene deletions that remove one α-globin copy (α^+^-thalassaemia), and (iii) non-deletional (ND) mutations that in some way inactivate the affected gene (α^ND^-thalassaemia). While deletions constitute the vast majority of these α-thalassaemia variants, non-deletional variants are typically associated with more severe phenotypes (10-12). Because the geographical distribution of β-thalassaemia largely overlaps with the distribution of α-thalassaemia, it is important to note that their co-inheritance often leads to a reduced imbalance between α-globin and β-globin chains, resulting in a milder thalassaemia phenotype (13-16).

From a clinical perspective, α-thalassaemia is mostly a burden in Southeast Asia where α^0^-thalassaemia variants (e.g. --^SEA^, --^THAI^) are common and result in HbH disease when inherited with α^+^-thalassaemia (e.g. -α^3.7^ or -α^4.2^) or α^ND^-thalassaemia (e.g. Hb Constant Spring, or Hb CS, or Hb Paksé), or in Hb Bart’s hydrops fetalis when inherited from both parents (17, 18). Previously, HbH disease was considered to be relatively benign; however, recent evidence suggests a spectrum of mild to severe forms of HbH disease, with the worst affected individuals requiring lifelong transfusion (10-12). Hb Bart’s hydrops fetalis, the most severe form of α-thalassaemia, associated with an absence of any functional α-globin genes, is almost always fatal *in utero* or soon after birth, although intrauterine interventions and perinatal intensive care can lead to survival (19).

In this context, there is a growing demand for a better understanding of the epidemiology of α-thalassaemia such that burden estimates can be calculated to guide public health decisions and assess the market for new pharmacological treatments. However, whilst several narrative reviews of the epidemiology of α-thalassaemia in Southeast Asian countries are available (18, 20, 21), a comprehensive review for the whole region has not been performed, making the current evidence-base patchy and incohesive. In addition, there appears to be a substantial amount of data that are available only in local data sources, which is not being accessed by the international community. Estimates of the number of newborns with severe forms of α-thalassaemia published by Modell and Darlison currently represent the only source of information on the epidemiology of thalassaemias and other inherited haemoglobin disorders at national and regional levels (22). Various inconsistencies have been identified in these estimates for α-thalassaemia (1). Furthermore, they do not include most of the surveys conducted in the genomic era, which has allowed accurate diagnosis through DNA testing. Finally, haemoglobinopathies often present remarkably heterogeneous geographical distributions (23, 24). As shown for other genetic conditions, these variations can be captured by generating continuous allele frequency maps interpolated from population surveys using geostatistical techniques. Combined with high-resolution demographic and birth rate data, these maps allow sub-national newborn estimates to be calculated (24, 25).

The aims of this study are therefore three-fold: i) to compile a geodatabase of published evidence for the distribution of α-thalassaemia and its common genetic variants in Southeast Asia, (ii) to generate the first model-based continuous maps of α-thalassaemia in Thailand and calculate refined estimates of the annual number of newborns affected by severe forms of α-thalassaemia, and iii) to comprehensively evaluate and summarise the compiled evidence-base for the whole region.

## Results

### The database

Our keyword searches yielded a total of 868 unique potential sources of data on α-thalassaemia prevalence and/or genetic diversity in 10 Southeast Asian countries: Brunei Darussalam, Cambodia, Indonesia, Lao People’s Democratic Republic, Malaysia, Myanmar, Philippines, Singapore, Thailand and Vietnam (Supplementary figure 1). A further 74 potential data sources were identified by one of the authors (SE) from local Thai journals and independently double-checked for inclusion (CH). Of all these sources, 75 met our inclusion criteria and were included in the final database. Due to some sources reporting estimates for more than one population, data were available for 106 individual population samples: 58 from the online literature search and 48 from the local literature. A detailed description of the database is provided in the Supplementary results.

**Figure 1.**
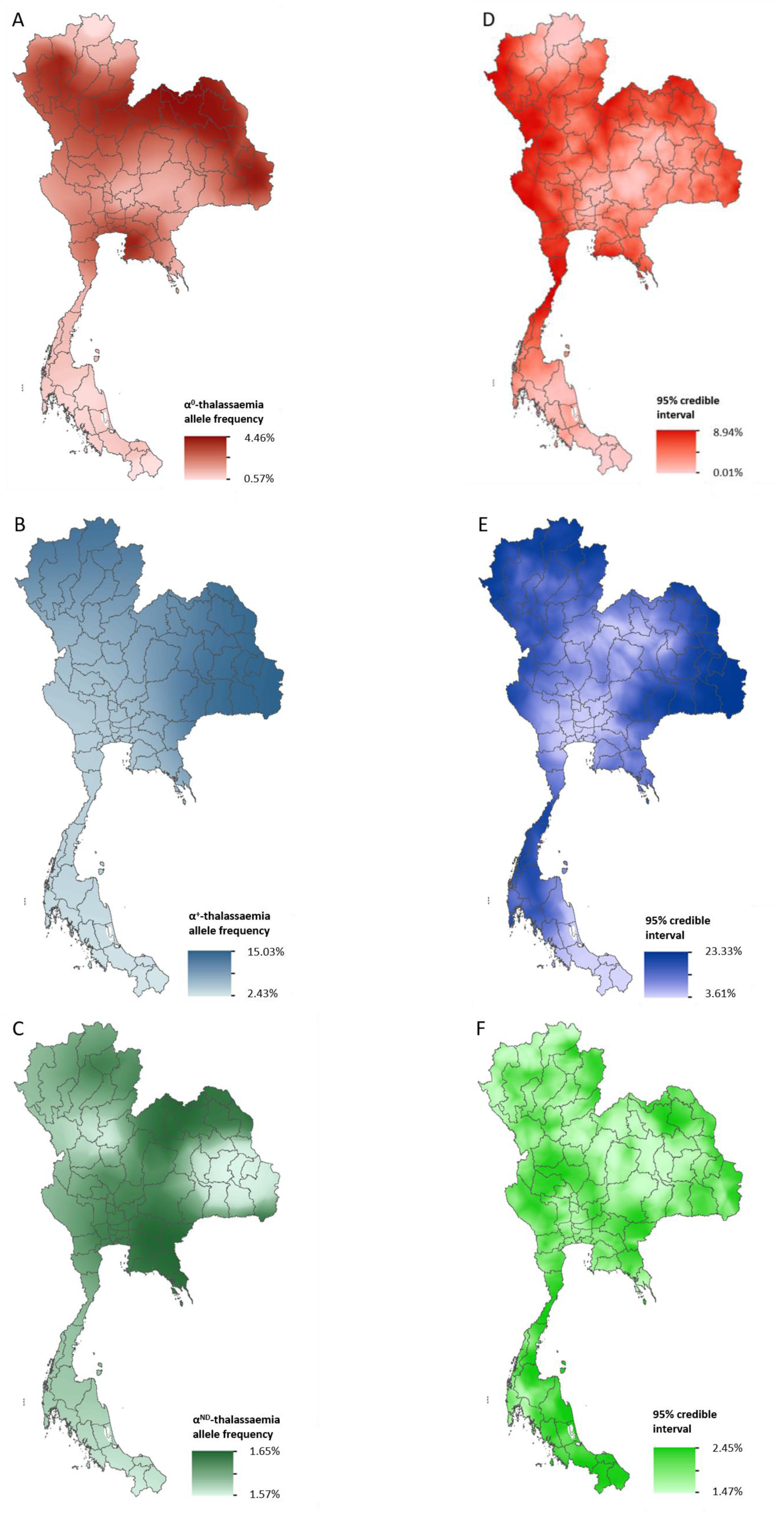
Maps of the mean of, and uncertainty in, the predicted α-thalassaemia allele frequencies in Thailand. Panels A to C display the mean of the posterior predictive distribution (PPD) of 100 realisations of the geostatistical model. Panels D to F display the 95% credible interval of the PPD. Each row corresponds to a different α-thalassaemia form: α^0^-thalassaemia (A and D); α^+^-thalassaemia (B and E) and α^ND^-thalassaemia (C and F).

Forty-six surveys provided data on α-thalassaemia gene frequency alone, two provided data only relating to genetic diversity and 58 provided data on both. Four surveys were reported at the national level (two from Malaysia, one from Singapore and one from Thailand), and were retained for the regional analysis. The spatial and temporal distributions of identified surveys are shown in Supplementary figure 2. The country for which the highest number of surveys was published in the international literature (i.e. excluding surveys obtained through local searches) was Thailand. No published surveys were identified for Brunei Darussalam or the Philippines. Within countries, surveys predominated in certain areas. For instance, in Thailand, the northern and northeastern parts of the country contained >75% of all surveys. Data for southern Thailand came exclusively from Thai journals (*n* = 4) (Supplementary figure 2A). The total number of individuals sampled was 133,649 (the population of the region is estimated to be 647,483,729 in 2017), with a mean sample size of 1,261.

**Figure 2.**
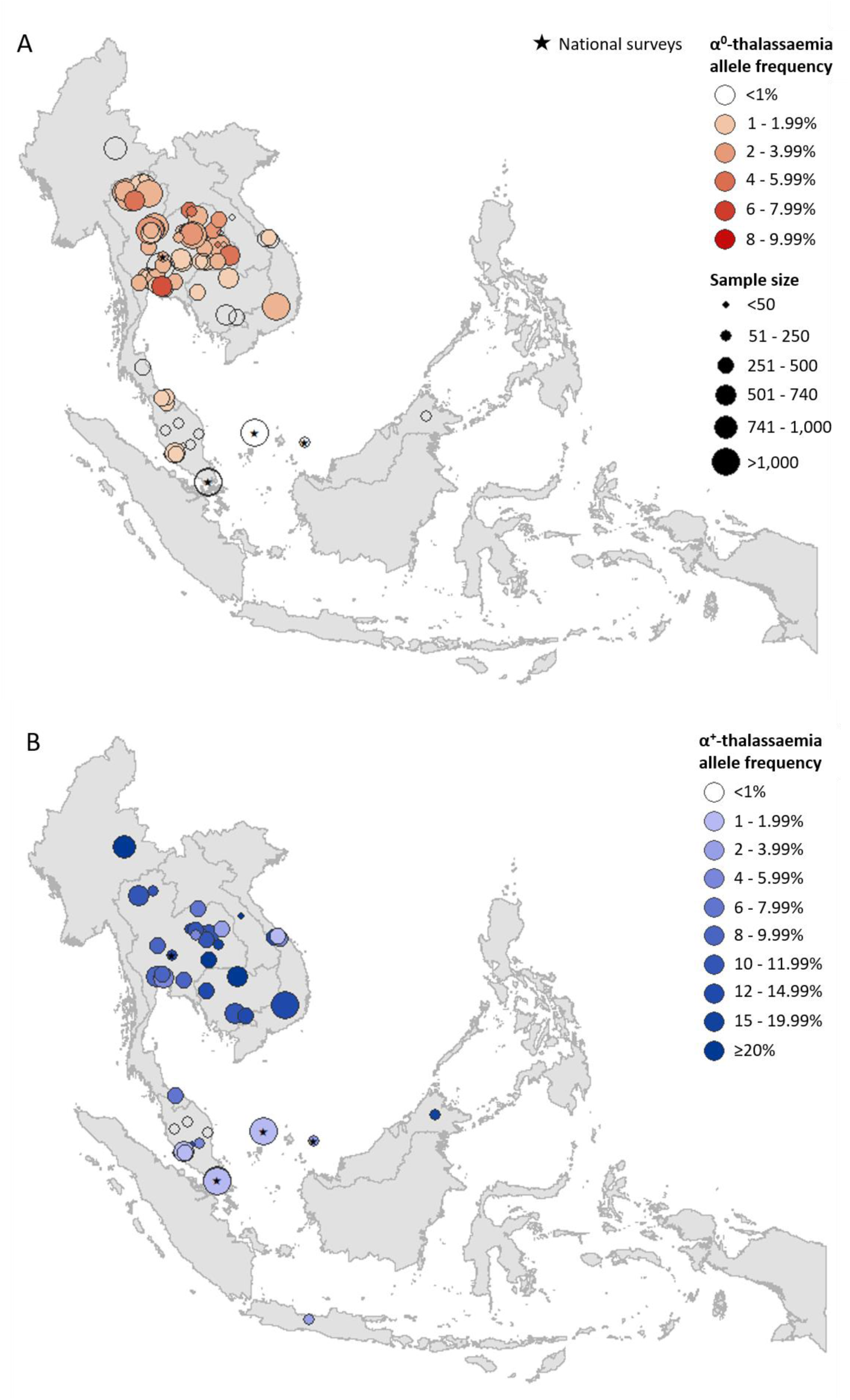
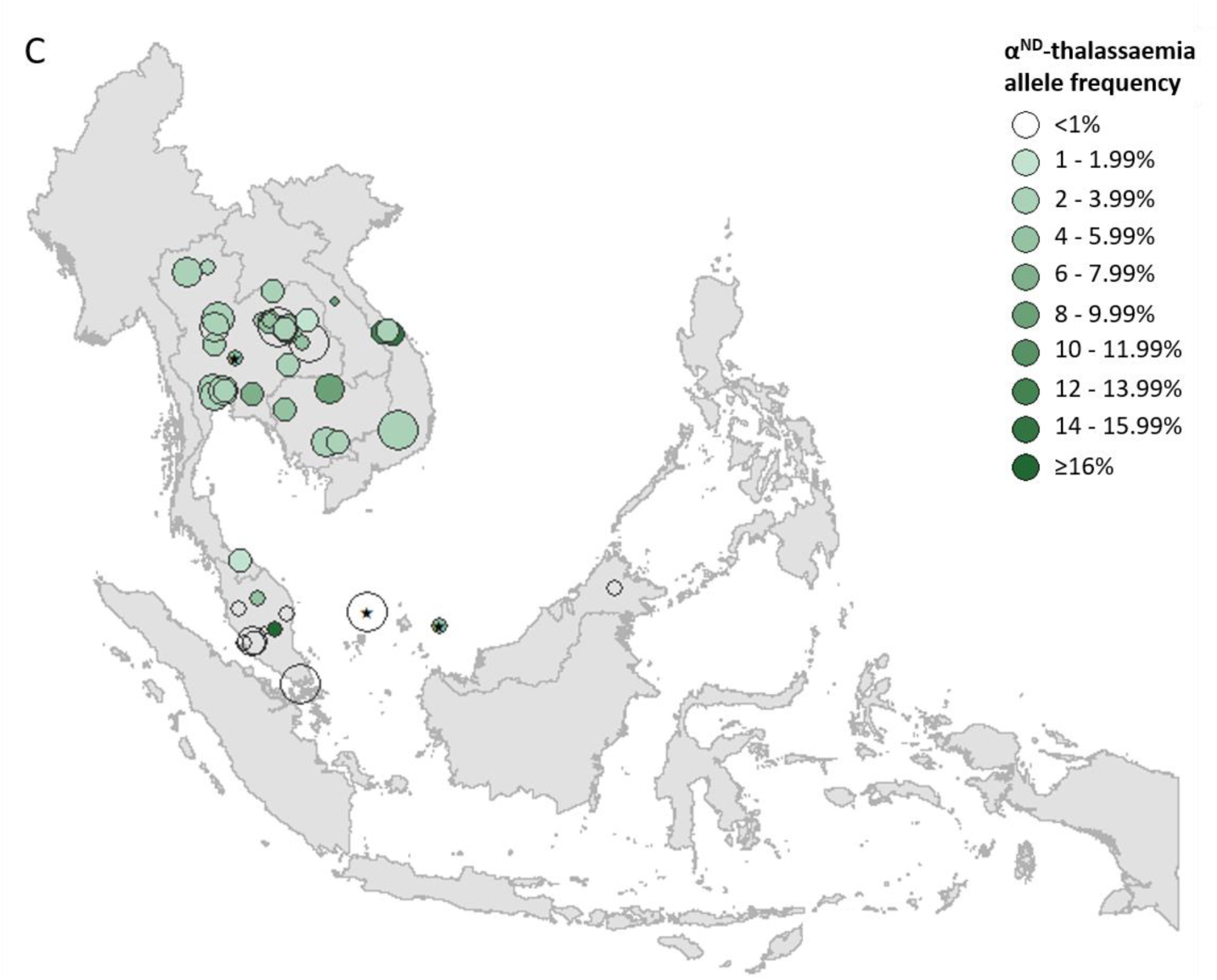
Descriptive maps of the observed allele frequencies in the database. (A) α^0^-thalassaemia, (B) α^+^-thalassaemia and (C) α^ND^-thalassaemia (on next page). A spatial jitter of up to 0.3^0^ latitude and longitude decimal degree coordinates was applied to allow visualisation of spatially duplicated data points. Colour intensity indicates allele frequency; circle size represents the size of the survey size.

Prevalence surveys varied considerably with regards to the α-thalassaemia alleles and/or genotypes that were tested for or reported upon; whilst some reported allele frequencies for α^0^-, α^+^- and α^ND^-thalassaemia, others provided data for only one or two of these. To maximise use of the available allele frequency data, whilst avoiding the incorporation of potentially biased estimates for overall α-thalassaemia allele frequency, we generated separate maps for each of the three major forms of α-thalassaemia – that is, α^0^-, α^+^- and α^ND^-thalassaemia (Figures 2A, B and C, respectively). α^0^-thalassaemia was the most extensively studied form (*n* = 97), followed by α^ND^-thalassaemia (*n* = 50) and then α^+^-thalassaemia (*n* – 47).

### Continuous allele frequency maps for Thailand

Data for Thailand and its neighbouring countries (Myanmar, Lao PDR, Cambodia and Malaysia) formed the evidence-base for a Bayesian geostatistical model. The total number of data points available for α^0^-, α^+^- and α^ND^-thalassaemia was 46, 39 and 41, respectively. The data were used to generate 1km x 1km maps of allele frequencies of α^0^-, α^+^- and α^ND^-thalassaemia in Thailand (Figure 1). One hundred realisations of the model were performed to generate a posterior predictive distribution (PPD) for each 1km x 1km pixel. The mean of the PPD is displayed, along with the 95% credible interval as a measure of model uncertainty.

The maps for α^0^- and α^+^-thalassaemia indicate clear spatial heterogeneity in allele frequencies, with ranges of 0.57%–4.46% and 2.43%–15.03%, respectively (Figure 1A, B). Heterogeneity is greatest in the north of the country. For α^0^-thalassaemia, while large parts of the northernmost provinces of Chiang Rai, Phayao and Nan have predicted allele frequencies of up to 2%, allele frequencies for the neighbouring provinces of Chiang Mai, Lampang and Phrae are often twice as high. The allele is also predicted at frequencies of up to 4% in the northeast of the country, along a belt across most of the north of the country and in Chonburi and Rayong provinces in central Thailand. Allele frequencies below 1% are predicted throughout the southern zone. α^+^-thalassaemia has its highest predicted allele frequencies across the whole of the north and northeastern zones. Predicted allele frequencies of α^ND^-thalassaemia range between 1.57% and 1.65% only.

Model uncertainty is greatest in areas where data are scarce (e.g. southern Thailand and along the border with Myanmar) or where there is heterogeneity in the available data (e.g. in Chiang Mai and the surrounding area). Overall, uncertainties are higher for α^+^-thalassaemia than for α^0^-thalssaemia or α^ND^-thalassaemia, which is partly due to the wider range of observed frequencies for this form. For α^0^-thalassaemia, the highest level of uncertainty is 9% and is found in Chumphon and Ranong provinces in southern Thailand and Kanchanaburi and Tak in the westernmost part of the country. For α^+^-thalassaemia, the highest uncertainty (up to 23%) is observed in the northeastern zone and in the north. Uncertainty for α^ND^-thalassaemia is patchy and ranged from 1.5% in central and northern Thailand to 2.5% in southern and northeastern Thailand. The results of the 10-fold cross-validation procedure reveal an average mean absolute error of the predictions of 0.93%, 4.10% and 2.30%, for α^0^-, α^+^- and α^ND^-thalassaemia, respectively. The average correlation between the observed and predicted values is 0.74 (0.62-0.83), 0.71 (0.49-0.85) and 0.47 (0.17-0.69), respectively.

### Estimates of number of affected newborns in Thailand

Estimates of the number of newborns born with a severe form of α-thalassaemia (i.e. Hb Bart’s hydrops fetalis and HbH disease) in Thailand in 2020 were generated by pairing our allele frequency predictions to high-resolution demographic data for the country. We estimate that the number of Hb Bart’s hydrops fetalis births in the country will be 423 (CI: 184 - 761) in 2020. The number of new cases of HbH disease is estimated to be 3,172, including 2,674 (CI: 1,296 - 4,491) deletional and 498 (CI: 237 - 947) non-deletional cases. The highest absolute burden of hydrops fetalis is predicted in Bangkok (57 [CI: 13 - 151) and Udon Thani (23 [CI: 6 - 66]) in the northeastern zone, despite the former not having the highest allele frequencies of α^0^-thalassaemia. Other states with a comparatively high burden include: Chiang Mai in the north of the country; Khon Kaen, Sakon Nakhon and Ubon Ratchathani in the northeast; and Chon Buri, Samut Prakan and Nonthaburi close to Bangkok. The estimated number of hydrops fetalis births in these provinces range between 10 and 19. For HbH disease, the highest burden is predicted in northeast Thailand for both the deletional and non-deletional forms. Bangkok is predicted to have the highest burden of HbH disease (301 [CI: 94 - 639] for deletional HbH disease and 68 [CI: 25 - 148] for non-deletional HbH disease).

To compare the above estimates with those previously published by Modell and Darlison, we also calculated our estimates using population and birth rate data for 2003 (Supplementary methods 3). Modell and Darlison estimated 1,017 and 2,515 births to be affected by Hb Bart’s hydrops fetalis and HbH disease, respectively, in 2003. Using population data from the same year paired with our model-based maps, and assuming no consanguinity, we estimate 709 and 5,469 newborns to be born with Hb Bart’s hydrops fetalis and HbH disease in the country. As Modell and Darlison included a population coefficient of consanguinity (*F*) in their calculations, we examined the effect that this would have on our estimates. We found that they do not change considerably (951 and 5,409), when a value of *F* of 0.01, a relatively high value for the region (www.consang.net), is incorporated. Our estimates are therefore consistent with those by Modell and Darlison for Hb Bart’s hydrops fetalis. However, they suggest that the burden of HbH disease in Thailand may have previously been underestimated. Moreover, whilst Modell and Darlison did not estimate the burden of non-deletional forms of HbH disease, our estimates suggest that 15% of the 5,469 neonatal cases were of non-deletional types, which are usually associated with more severe phenotypes.

### Overall distribution of α-thalassaemia across Southeast Asia

In our database for all of Southeast Asia, the number of surveys that tested for all three forms of α-thalassaemia was 40. Amongst these, the overall α-thalassaemia gene frequency ranged from 0% in populations from peninsular Malaysia to 35.4% in Preah Vihar, Cambodia (26). A higher allele frequency of 49% was also reported in the So ethnic group from Khammouane Province in Lao PDR, although the sample size for this study was small (*N* = 50) (27). Supplementary table 2 shows the observed allele frequency ranges for the different α-thalassaemia forms in each country.

For α^0^-thalassaemia, the highest allele frequencies were observed in Thailand (Figure 2A) In Lao PDR, surveys along the Lao PDR-Thailand border near Vientiane reported allele frequencies between 4.03% and 7.28%, whilst the survey among the aforementioned So ethnic group reported an absence of the α^0^-thalassaemia allele. The highest reported allele frequencies in Cambodia and Vietnam were 1.10% and 2.66%, respectively, with the majority of studies reporting even lower frequencies. However, data were sparse in the two countries (*n* = 4 in each). Allele frequencies of up to 1.53% were observed in southern Thailand, whilst the few surveys carried out in Myanmar (*n* = 1), Malaysia (*n* = 11) and Singapore (*n* = 2) reported allele frequencies of around 1%. In Malaysia, the highest allele frequency of α^0^-thalassaemia was 1.92% from a study carried out in newborns in Kuala Lumpur, the capital city (28).

α^+^-thalassaemia, the most prevalent form, reached allele frequencies of 26% in Cambodia (Figure 2B) (26). The surveys revealed a clear north-to-south decline in the distribution of α^+^-thalassaemia across the region, with a single high allele frequency estimate of 16.8% observed in Sabah in Malaysian Borneo (29). High allele frequencies (≥10%) were observed in all four surveys in Cambodia. In Vietnam the reported α^+^-thalassaemia allele frequency ranged from 1.59% to 14.4%.

The observed allele frequency of α^ND^-thalassaemia ranged between 0% in various locations across Malaysia and 16.25% in central Peninsular Malaysia (Figure 2C). Within Thailand, the highest reported allele frequencies of around 7% were observed in Khon Kaen in the northeast and Chachoengsao in central Thailand. In Cambodia and Vietnam, α^ND^-thalassaemia allele frequencies of up to 8% and 14.3% were reported, respectively, in surveys in which α^0^-thalassaemia was found to be absent, whilst in other parts of these countries, the two forms were found to co-exist at similar allele frequencies (e.g. around 2.5% in Binh Phuoc and Khanh Hoa provinces in Vietnam).

### Genetic diversity of α-thalassaemia across Southeast Asia

Maps of the genetic diversity of α-thalassaemia across Southeast Asia are shown in Figures 3-6. Figure 3 displays surveys that included all three α-thalassaemia forms (α^0^-, α^+^- and α^ND^-thalassaemia) and only distinguished between them. Figures 4-6 display surveys that provided information on the frequencies of specific α-thalassaemia variants (e.g. --^SEA^, -α^3.7^, etc.). For these, the variants that were tested for differs between surveys. Some surveys are included in both Figure 3 and Figures 4-6. For the latter figures, the region has been divided to improve visualisation of the data.

**Figure 3.**
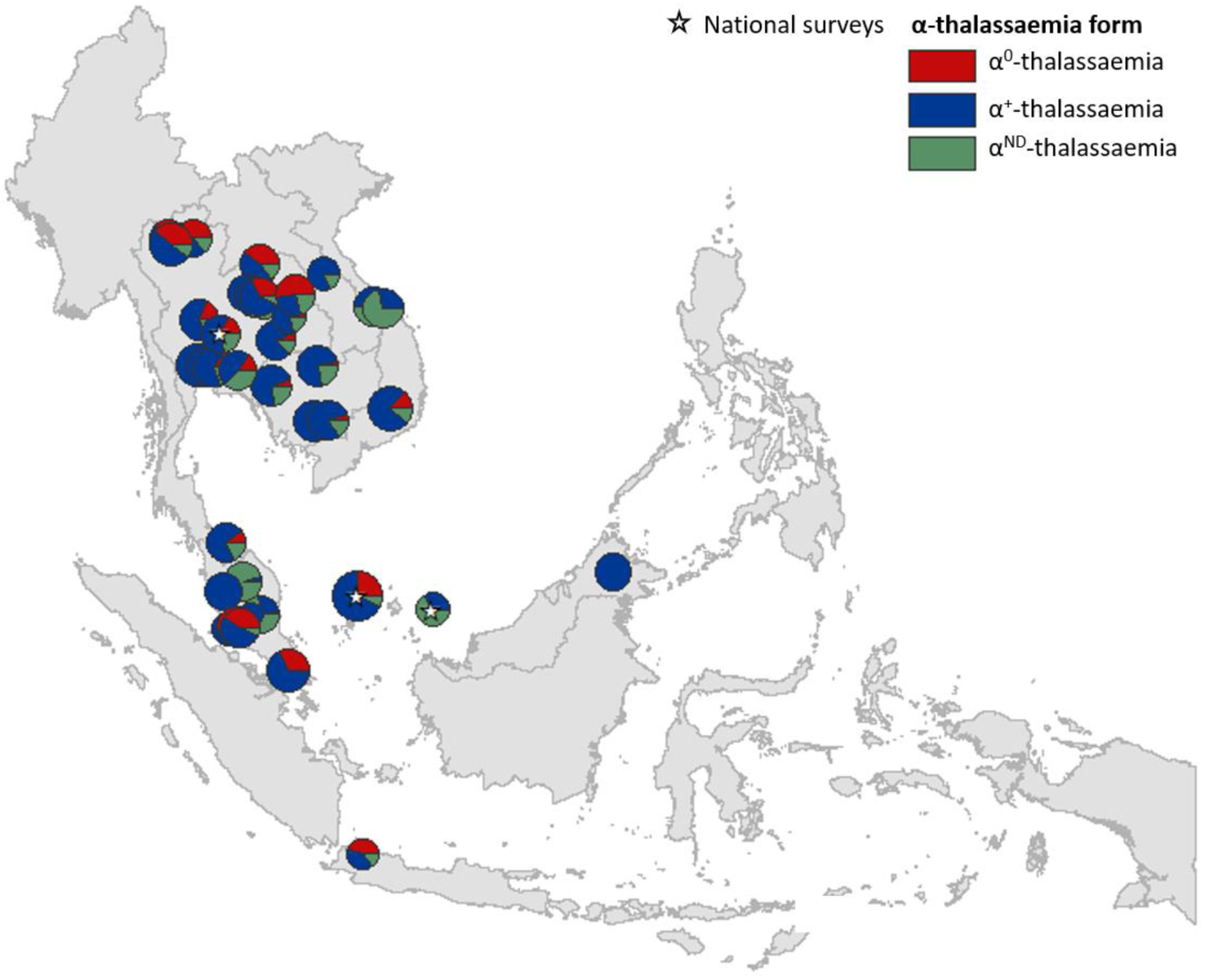
Map showing the proportions of α^0^-, α^+^- and α^ND^-thalassaemia in Southeast Asia. Three surveys were mapped at the national level (indicated by a white star). The size of the pie charts reflects survey size on a log scale.

**Figure 4.**
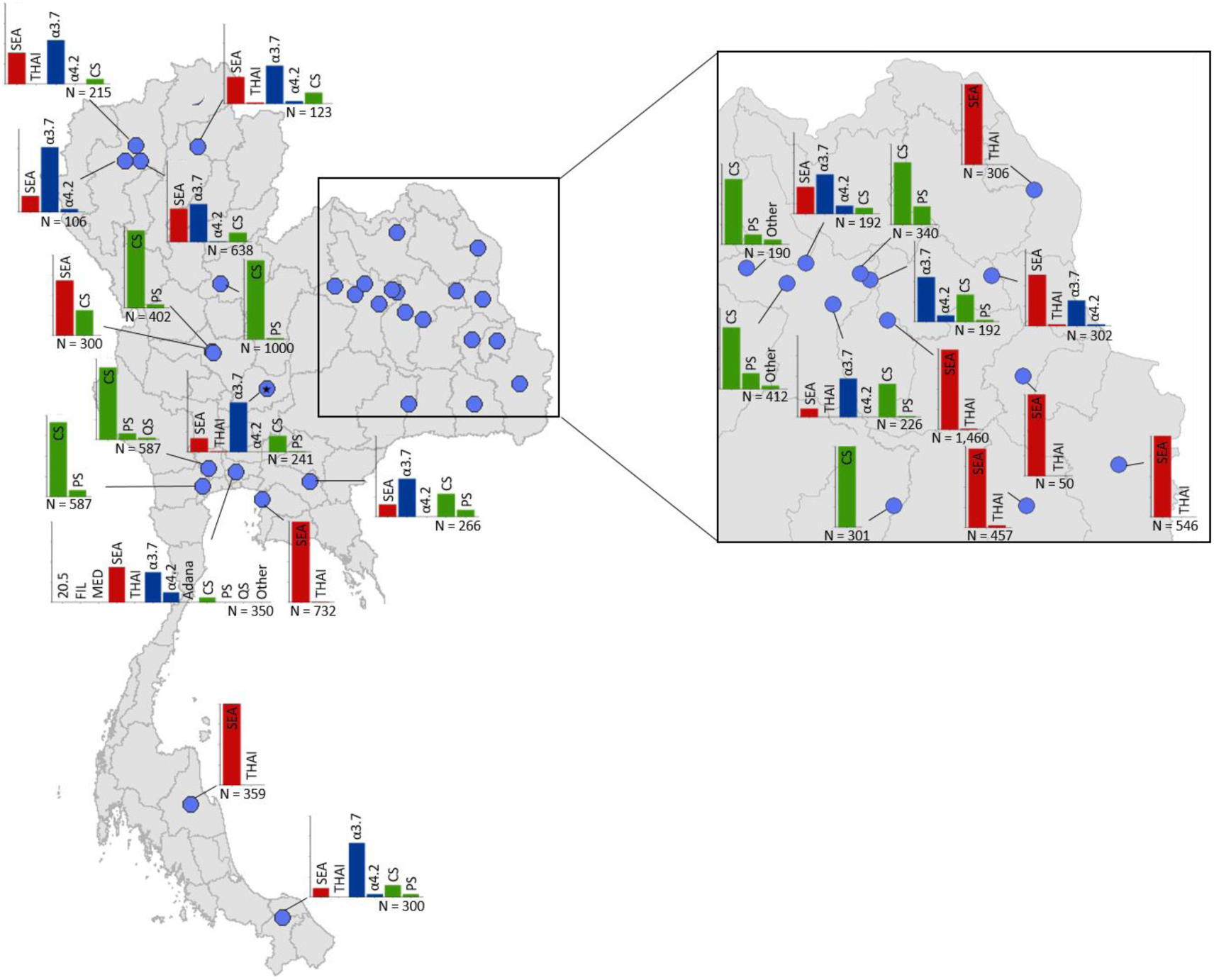
Map showing the proportions of specific α-thalassaemia variants in Thailand. Given the high number of surveys in northeast Thailand, this region has been magnified. The *y*-axis scale is the same across all bar charts, ranging from 0 to 1. The variants that were tested for in each survey are indicated above each bar. α^0^-thalassaemia mutations are shown in red, α^+^-thalassaemia mutations in blue and α^ND^-thalassaemia mutations in green. Empty spaces along the *x*-axis indicate an absence of the corresponding mutation in the survey sample. The sample size of the survey is given under each plot. Bar charts are connected to their spatial location by a black line.

**Figure 5.**
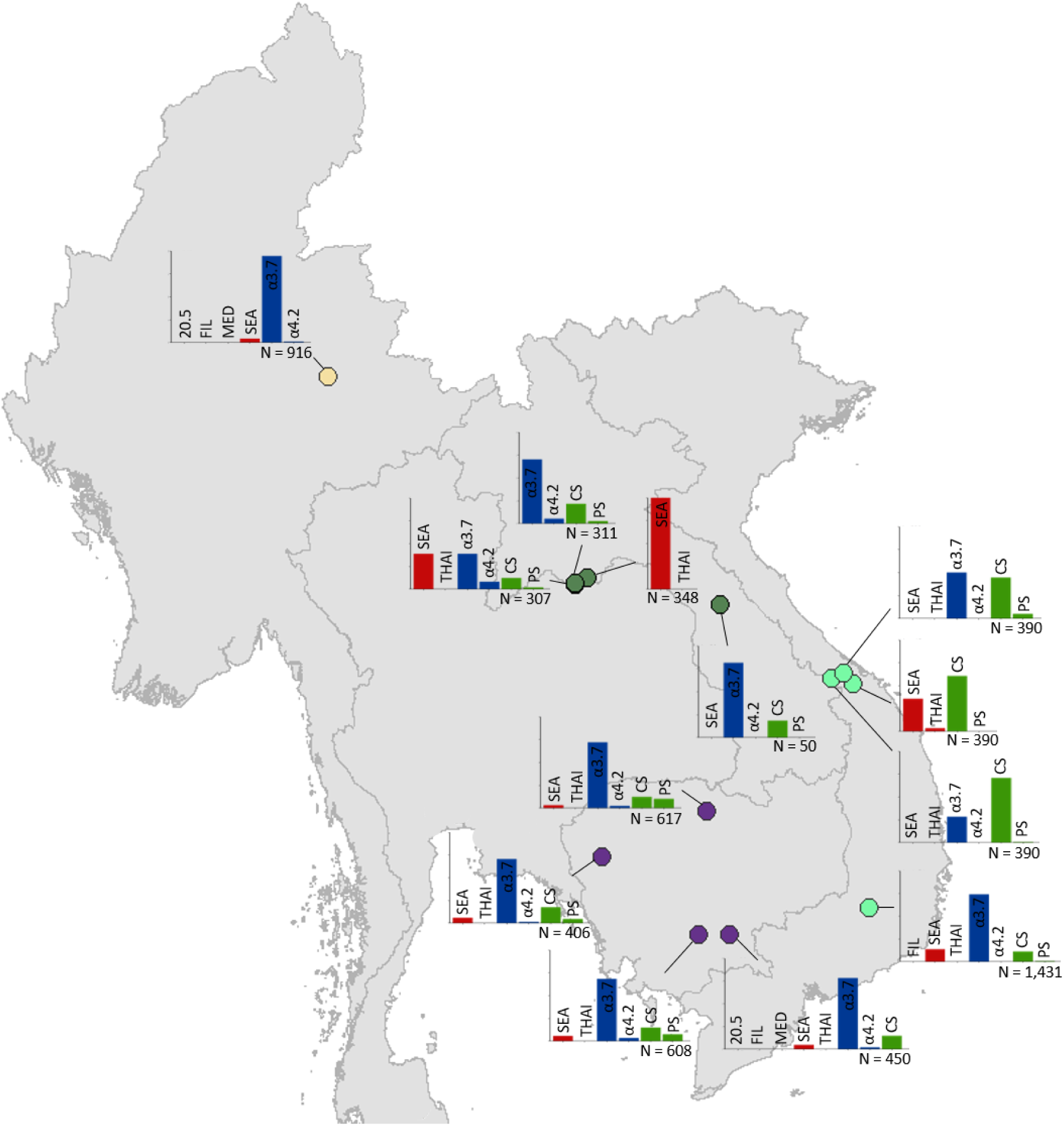
Map showing the proportions of specific α-thalassaemia variants in Myanmar, Lao PDR, Cambodia and Vietnam. The *y*-axis scale is the same across all bar charts, ranging from 0 to 1. The variants that were tested for in each survey are indicated above each bar. α^0^-thalassaemia mutations are shown in red, α^+^-thalassaemia mutations in blue and α^ND^-thalassaemia mutations in green. Empty spaces along the *x*-axis indicate an absence of the corresponding mutation in the survey sample. The sample size of the survey is given under each plot. Bar charts are connected to their spatial location by a black line. Data points are coloured by country, using the same colour scale as that in Supplementary Figure 1.

**Figure 6.**
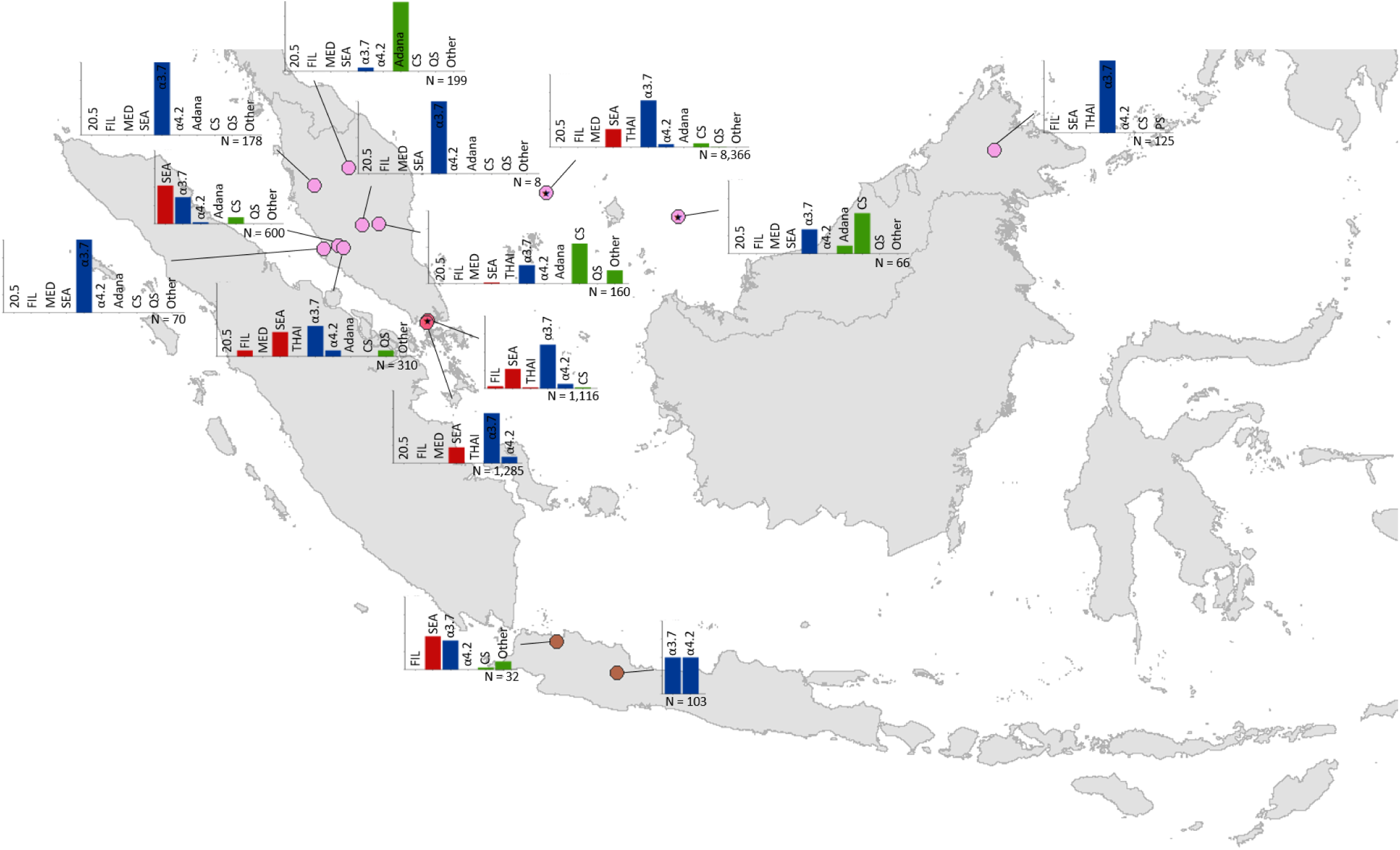
Map showing the proportions of specific α-thalassaemia variants in Malaysia, Singapore and Indonesia. The *y*-axis scale is the same across all bar charts, ranging from 0 to 1. The variants that were tested for in each survey are indicated above each bar. α^0^-thalassaemia mutations are shown in red, α^+^-thalassaemia mutations in blue and α^ND^-thalassaemia mutations in green. Empty spaces along the *x*-axis indicate an absence of the corresponding mutation in the survey sample. The sample size of the survey is given under each plot. Bar charts are connected to their spatial location by a black line. Data points are coloured by country, using the same colour scale as that in Supplementary figure 1.

α^+^-thalassaemia most consistently constituted the highest proportion of α-thalassaemia, although there were some surveys in which α^ND^-thalassaemia was the predominant form (e.g. central Vietnam and in parts of Malaysia). In Figure 3, areas where the observed relative proportion of α^0^-thalassaemia was greatest include: Chiang Mai and Phayao provinces in north Thailand, Kalasin in northeast Thailand, Vientiane in Lao PDR, Kuala Lumpur and Selangor in Malaysia, Singapore and Jakarta in Indonesia. The α^0^-thalassaemia allele was absent in the survey from Malaysian Borneo as well as in central Vietnam and central Lao PDR. In certain areas, α^0^- and α^ND^-thalassaemia together accounted for the majority of α-thalassaemia (e.g. ∼75% in Kalasin in Thailand, ∼60% in Kuala Lumpur and Jakarta and ∼53% in Khon Kaen and Chachoensao in Thailand and Vientiane in Lao PDR). Some of these areas also correspond to where the highest allele frequencies of these alleles are found, for example, northeast Thailand and the Thailand-Lao PDR border.

Among surveys that tested for specific α-thalassaemia variants, the most commonly tested variant throughout the region was --^SEA^ (*n* = 44), followed by -α^3.7^ (*n* = 36) (Figures 4-6). Most of the studies in Thailand tested for a subset of the mutations considered in this study; only one survey in Bangkok tested for the whole suite. More than in other countries, surveys in Thailand tested specifically for α^0^- or α^ND^-thalassaemia mutations (*n* = 7 and 9, respectively).

Throughout the region, --^SEA^ was the dominant α^0^-thalassaemia mutation, and in the majority of surveys -α^3.7^ was the dominant α^+^-thalassaemia and Hb CS the dominant α^ND^-thalassaemia mutation. The only exceptions were in Java in Indonesia, where -α^3.7^ and - α^4.2^ were found in equal proportions and in Kelantan in Malaysia, where Hb Adana was the only α^ND^-thalassaemia mutation identified. The -(α)^20.5^ and --^MED^ mutations were not detected in any of the surveys, whilst the --^FIL^ mutation was found in 2 of the 16 surveys in which it was tested for and --^THAI^ in 9 of the 31 surveys in which it was included. Consistent with Figure 3, α^0^-thalassaemia variants accounted for a small proportion of α-thalassaemia mutations in Myanmar, Cambodia and Vietnam and a high proportion in surveys along the Thailand-Lao PDR border. In Malaysia, Indonesia and Singapore, the proportion of α^0^-thalassaemia varied considerably, with it being absent in some areas and the predominant form in others. This is also true for α^ND^-thalassaemia.

## Discussion

α-thalassaemia is a neglected public health problem whose burden has, to date, been largely overlooked, but for which morbidity is expected to increase in the coming decades as a result of the epidemiological transition (1, 30). Moreover, country reports (e.g. from Malaysia) indicate a shift in the age distribution of thalassaemia patients towards older ages (31). As the burden increases, there will be greater demand for resources, including healthcare facilities and staff, genetic counselling and drugs, to treat and manage affected patients. This is particularly true for countries in Southeast Asia, where severe forms of α-thalassaemia are found at high prevalence.

### Comparison with existing maps and population estimates

The model-based maps for Thailand presented here are, to our knowledge, the first spatially continuous maps of the distribution of α-thalassaemia in any country. Our newborn estimates represent the first evidence-based estimates of specific forms of α-thalassaemia disease amongst newborns since 2003 (although the study in which they were reported was published in 2008)(22) and the first estimates at sub-national level. Importantly, whilst there are currently no estimates of the number of stillbirths that will occur in Thailand in 2020, our estimate of the number of Hb Bart’s hydrops fetalis births represents more than 10% of the number of stillbirths estimated for 2015 (3,697) (32).

Comparisons between previous newborn estimates and those generated in this study using our updated database and 2003 demographic data revealed an almost two-fold difference for deletional HbH disease (2,515 compared to 4,694 in the present study). Reasons for such discrepancies most likely relate to: (i) differences in the inclusion criteria used in the generation of our map and therefore our calculation of newborn estimates, (ii) the quantity of survey data used, and iii) the statistical methods employed. For instance, spatial specificity was not a consideration in the study by Modell and Darlison, who used a single allele frequency estimate extrapolated to the whole country. As such, the newborn calculations in the present study represent a methodological advance over previous efforts to assess the burden of α-thalassaemia. We related fine-scale allele frequency data to birth count data of equally high resolution, allowing location-specific estimates to be generated that could be aggregated to province level. Moreover, the use of model-based maps in our calculations enabled the measurement of uncertainty in our predictions. Finally, by including allele frequency data on α^ND^-thalassaemia, we were able to estimate the burden of the more severe non-deletional HbH disease.

Our newborn estimates for 2020 are considerably lower than those for 2003. This reduction is due to the lower number of births in Thailand in 2020 as a result of a decreasing birth rate and population size (33). It would be interesting to quantify how improvements in the prevention of thalassaemias will affect these estimates in the future.

Our descriptive maps represent the first detailed cartographic representations of α-thalassaemia allele frequency estimates in Southeast Asia, which take into account the specific geographical location of the surveys in which they were observed. Until now, available maps (e.g. Supplementary figure 3) provided only a crude overview of overall α-thalassaemia gene frequency, without any distinction between different α-thalassaemia forms, and extrapolated to the entire region, thereby masking sub-national and even international, variation in allele frequencies (1).

The maps are broadly consistent with early narrative reviews of the gene frequency of α-thalassaemia in the region (20), showing a clear north-to-south trend of decreasing allele frequencies of α^0^- and α^+^-thalassaemia and a patchier distribution of α^ND^-thalassaemia. However, our maps also demonstrate a severe lack of data on the allele frequency of α-thalassaemia across large parts of Southeast Asia, including in Myanmar, northern Lao PDR, northern Vietnam, Indonesia, Philippines and Brunei. This impedes our ability to assess the fine-scale burden of α-thalassaemia, making efficient public health planning for its control difficult, and limits our ability to track progress in the prevention and management of the disorder.

### Public health implications of patterns of genetic variation

The pattern of genetic diversity observed in this study indicates variable distributions of mild and severe α-thalassaemia forms. This has important implications for the design of newborn screening programmes with regards to the preferred diagnostic algorithm and allocation of treatment and management service provision. Areas with the highest proportions of co-occurring severe α-thalassaemia forms (i.e. α^0^-thalassaemia and α^ND^-thalassaemia) may experience a higher prevalence of the severe non-deletional form of HbH disease. Furthermore, the predominance of Hb CS in surveys from Malaysia and Vietnam suggests that the health burden of α-thalassaemia in these areas may be greater than previously thought. Hb CS is a mutation at the termination codon of the α2-globin gene, which, in a normal individual, accounts for three-quarters of overall α-globin production (34, 35). As a result, α2-globin gene mutations, such as Hb CS, tend to cause a more severe phenotype (18).

### Model strengths and limitations

The reliability of the model-based predicted maps is intrinsically linked to the quality, quantity and spatial coverage of the data upon which the models are based. We were unable to generate continuous maps for the whole of the Southeast Asian region as data were sparse in large areas. We are aware that unpublished surveys are likely to be available for most countries of the region. We showed that a substantial additional amount of survey data could be identified in local published sources in Thailand. Obtaining local data for all of the countries in southeast Asia was beyond the scope of this study, but what was possible for Thailand shows the enormous value of future collaborations to collate local data in other regions.

For Thailand, limitations relating to data sparsity, uneven survey distributions and allele frequency heterogeneity can be quantified in the presented uncertainty intervals. Areas where there is little data or where observed allele frequencies are highly heterogeneous within a small geographical area will have more uncertain predictions, whilst a large amount of data for which there is little heterogeneity will lead to more precise predictions. We identified a lack of data in the southern part of Thailand, which is reflected in larger uncertainty estimates. Other predictions with high associated uncertainty include those along the Thailand-Myanmar border, where no surveys on α^0^-thalassaemia prevalence are found. This highlights the arbitrary nature of country borders in mapping studies.

Extensive variation in allele frequencies between different ethnic groups has been observed (36). Due to the nature of this study, and the smoothing of allele frequencies over a continuous spatial domain, our predicted allele frequencies might not fully reflect heterogeneity between local ethnic populations. For example, allele frequencies of around 3.65% for the Hb CS mutation have been reported in the Khmer ethnic group in Surin and Buriram provinces, whilst our model predicts maximum allele frequencies of 1.65% here. It is likely that other factors influence the allele frequencies of the different α-thalassaemia forms, which have not been considered in this mapping study, including ethnicity, consanguinity, rates of malaria (both *Plasmodium falciparum* and *P. vivax*)(37) and population migration patterns. Furthermore, there is bound to be uncertainty in the geolocation of some of the surveys included in the study due to the lack of details published or available. This uncertainty could not be accounted for. Finally, whilst the inclusion criterion of molecular methods should help to improve the reliability of allele frequency estimates, they are not 100% sensitive(38) and do not cover all possible α-thalassaemia mutations, which may lead to some error in the reported allele frequencies.

Whilst we have calculated the burden of α-thalassaemia in terms of the number newborns born with severe forms in 2020, there are other aspects of the disease burden that would be worth considering pending the availability of more data, for example, milder-forms and their coinheritance with β-thalassaemia, DALY losses from α-thalassaemia, maternal complications (some of which can be life-threatening) (18, 39), psychological effects and, in the case of HbH disease, survival data allowing the calculation of all-age population estimates. Furthermore, the estimates presented here do not include compound disorders, such as EA Bart’s and EF Bart’s diseases (HbH disease with heterozygous and homozygous forms of β^E^, another clinically important structural β-globin variant, respectively) (40). Finally, the visualisation of our burden estimates are subject to the modifiable area unit problem, whereby the presentation of estimates at the province level likely masks pockets of high burden (41).

### Future prospects and conclusions

The allele frequency, distribution and genetic variant profile of α-globin forms is only a part of their epidemiological complexity. An improved understanding of the natural history of α-thalassaemia and the factors that modify its clinical outcome will be imperative for establishing better estimates of its burden. This is particularly pertinent in the Southeast Asian region, where the disorder co-exists with β-thalassaemia, including the commonest haemoglobin variant, Hb E. Many studies have shown a positive epistatic interaction between α- and β-thalassaemia, whereby their co-inheritance results in the amelioration of the associated blood disorder (42, 43).

A detailed assessment of current knowledge on the allele frequency of α-thalassaemia and the magnitude of its health burden is needed to develop suitable prevention and control programmes. This study provides a detailed overview of the existing data on the gene frequency and genetic diversity of α-thalassaemia in Southeast Asia. We show that our knowledge of the accurate allele frequency and distribution of this highly complex disease remains somewhat limited. Because of the remarkable geographic heterogeneities in the gene frequency of α-thalassemia, interventions have to be tailored to the specific characteristics of the local population (e.g., prevalence of the disorder in the population, ethnic makeup, and consanguinity) and the local health care system As the epidemiological transition in these countries continues (30, 44, 45), it will become increasingly important to regularly update regional and national maps of α-thalassaemia gene frequency and newborn estimates such that health and demographic changes can be properly quantified (1). Our findings provide a baseline for such endeavours.

## Methods

### Compiling a geodatabase of α-thalassaemia allele frequency and genetic diversity

A comprehensive search of three major online bibliographic databases (PubMed, ISI Web of Knowledge and Scopus) was performed to identify published surveys of α-thalassaemia prevalence and/or genetic diversity in Southeast Asia (Figure 7). The 10 member states of the Association of Southeast Asian Nations (ASEAN) were used to define the region under study and include: Brunei Darussalam, Cambodia, Indonesia, Lao PDR, Malaysia, Myanmar, Philippines, Singapore, Thailand and Vietnam (Supplementary figure 1). In addition, for Thailand, articles published in national journals (in Thai) – not included in international bibliographic databases – were manually searched for local surveys. Consistent and pre-defined sets of inclusion criteria for prevalence/allele frequency data and genetic diversity data, outlined in the Supplementary methods 1, were used to identify relevant surveys. Data extracted from Thai journals were independently validated against the inclusion criteria by two of the authors (SE and CH). The assembled datasets are available in the Supplementary material.

**Figure 7.**
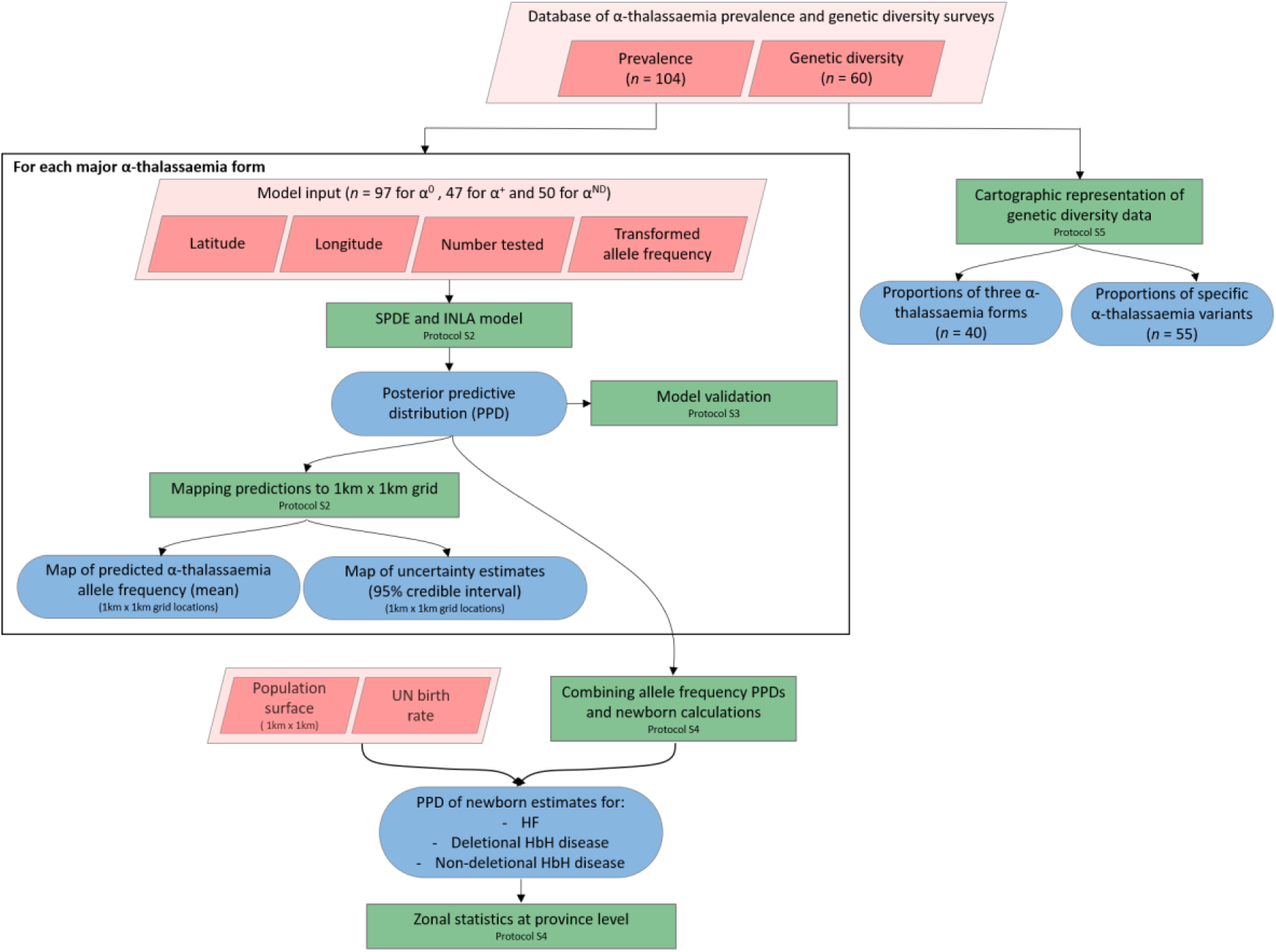
A schematic overview of the methodology used in this study and a breakdown of the data types analysed. Pink diamonds indicate the database and input data; green boxes denote model processes and data visualisation steps; blue rods represent study outputs.

### Modelling continuous maps of α-thalassaemia allele frequency in Thailand

We employed a Bayesian geostatistical framework to model the allele frequencies of α^0^-, α^+^- and α^ND^-thalassaemia, respectively, in Thailand, where a substantially higher number of surveys were identified. We included data from Thailand and its neighbouring countries (Myanmar, Lao PDR, Cambodia and Malaysia) in order to preclude the possibility of a border effect. Three surveys that were reported only at the national level (one in Thailand and two in Malaysia) were excluded for this part of the analysis. Only geographical location was included as a predictor of α-thalassaemia allele frequency (Figure 7).

For each of the three main forms of α-thalassaemia, a model was fitted using a Bayesian Stochastic Partial Differential Equation (SPDE) approach with Integrated Nested Laplace Approximation (INLA) algorithms developed by Rue *et al*. (46), available in an R-package (www.r-inla.org). The fitted model was then used to generate predictions at a resolution of 1km x 1km for α-thalassaemia allele frequencies for all unsampled locations in Thailand. Uncertainty estimates, measured as the 95% credible interval, for the predictions were calculated using 100 conditionally simulated realisations of the model to generate a posterior predictive distribution (PPD) for each 1km x 1km pixel. Full details of the modelling process and model validation procedures, which involved a 10-fold cross validation, are provided in the Supplementary methods 2.

### Refining estimates of the annual number of neonates affected by severe disease forms

To generate estimates of the annual number of newborns affected by Hb Bart’s hydrops fetalis syndrome (--/--) and deletional and non-deletional HbH disease (-α/-- and αα^ND^/--, respectively) in Thailand in 2020, we paired the predicted allele frequency maps generated using our Bayesian geostatistical framework with high-resolution birth count data. First, we combined the three allele frequency maps to estimate the frequency of each genotype in each pixel, assuming Hardy-Weinberg proportions for a four-allele system (Equation 1 and Table 1) (47, 48).

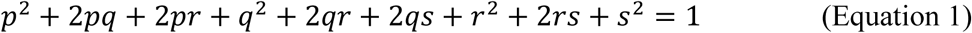

where, *p* is the allele frequency of α a^0^-thalassaemia (--), *q* is the allele frequency of α^+^-thalassaemia (-α), *r* is the allele frequency of α^ND^-thalassaemia (αα^ND^) and *s* is the allele frequency of the wild-type α-globin haplotype (αα).

**Table 1.** A breakdown of the genotypes for the three clinically important forms of α-thalassaemia – Hb Bart’s hydrops fetalis, deletional HbH disease and non-deletional HbH disease – and the Hardy-Weinberg equilibrium (HWE) proportions used for their calculation. To compare our model output with previous newborn estimates for Hb Bart’s hydrops fetalis and deletional HbH disease, we paired our allele frequency maps with 2003 demographic and birth data and included a measure of consanguinity in our calculations.

| Genotype | Disorder | HWE proportions | Inclusion of population coefficient of consanguinity (F) |
| --- | --- | --- | --- |
| --/-- | Hb Bart’s hydrops fetalis | $p^2$ | $p^2 + Fp(1 - p)$ |
| $-\alpha/--$ | Deletional HbH disease | $2pq$ | $2pq(1 - F)$ |
| $\alpha\alpha^{\text{ND}}/--$ | Non-deletional HbH disease | $2pr$ | $2pr(1 - F)$ |

To calculate birth counts, the 2015-2020 crude birth rate for Thailand was downloaded from the 2017 United Nations (UN) world population prospects(33) and multiplied with a high-resolution predicted 2020 population surface, adjusted to UN population estimates, obtained from the WorldPop project (www.worldpop.org.uk, last accessed 23 January 2018) (49). The predicted genotype frequencies were then paired with the birth count data over 100 conditionally simulated realisations of the geostatistical model and areal estimates at province level calculated, together with 95% credible intervals; their calculation is described in Supplementary methods 3. We also applied our maps to 2003 demographic data, and incorporated consanguinity into our calculations (Table 1), in order to more directly compare estimates generated using our method with previous estimates (22).

### Summarising the current evidence-base for α-thalassaemia gene frequency in Southeast Asia

Cartographic representations of the identified prevalence surveys were generated using ArcGIS 10.4.1 (ESRI Inc., Redlands, CA, USA). The descriptive maps reflect the spatial distribution of the prevalence surveys, along with their respective sample sizes and observed α^0^-, α^+^- and α^ND^-thalassaemia allele frequencies. Other features of the database, including the temporal distribution of the surveys, the identity of the populations studied (e.g. community, pregnant women, newborns, etc.) and the contribution of local Thai surveys to the evidence-base, were also examined. In addition, pairwise univariate linear regression was performed to examine the association between the allele frequencies of the different forms of α-thalassaemia in the dataset, transformed on a log scale.

### Mapping α-thalassaemia genetic diversity

Maps of the genetic diversity of α-thalassaemia across Southeast Asia were also generated (Figure 6). Given the heterogeneity in the reporting of different α-thalassaemia genotypes, we divided the genetic diversity data into two subtypes: (i) those surveys that only distinguished between the different α-thalassaemia forms (α^0^-, α^+^-, and α^ND^-thalassaemia), and (ii) those surveys that contained detailed count data for a range of common mutations. We focused on the 11 mutations that are most commonly reported in Southeast Asia or are part of standard multiplex polymerase chain reaction (PCR) methods: -α^3.7^, -α^4.2^, --^SEA^, --^THAI^, --^MED^, --^FIL^, - (α)^20.5^, Hb Adana (HBA2:c.179G > A), Hb CS (HBA2:c.427T > C). Hb Paksé (HBA2:c.429A > T), Hb Quong Sze (HbA2:c.377T > C) (50). An “Other” category was used for other, rarer α-thalassaemia mutations. For the first data subtype, only surveys that tested for all three α-thalassaemia forms were mapped and the relative proportions of the different forms in the study sample were displayed using pie charts. For the latter, the same approach to that used in Howes *et al.* (2013) was used; the variant proportions were displayed using bar charts in which all variants that were explicitly tested for were included on the *x*-axis (25). This allowed information regarding the suite of variants that were tested for in the survey to be displayed as well as unambiguous representation of the absence of a variant in the study sample.

## Author contributions

CH, SE, VV and FBP developed the conceptual approach. CH, SE and FBP assembled and abstracted the data. CH and SB implemented the modelling and computational tasks. CH, SB, BSP and FBP analysed the data. CH wrote the first draft of the report and generated figures. All authors contributed to the study design and data interpretation and to the revision of the final manuscript.

## Supplementary methods 1: Constructing a geodatabase of α-thalassaemia prevalence and/or genetic diversity surveys

### Library assembly

Three online biomedical literature databases (PubMed, ISI Web of Knowledge and Scopus) were systematically searched for all records referring to “α-thalassaemia”, “alpha-thalassaemia”, “α-thal” or alpha-thal”. Broad search terms were used to make the search as inclusive as possible. Retrieved articles were imported into the bibliographic database, Endnote X7 (Thomson Reuters, Carlsbad, CA, USA) and grouped by country. The countries included in this study are shown in Supplementary figure 1. The last search was conducted on 21 July 2017. In addition, a non-systematic review of surveys available in local Thai journals was performed.

Supplementary figure 1 here.

### Survey inclusion criteria

The title and abstract of each reference was reviewed for its suitability to this study and those that were considered not to be so were excluded from further review. This included animal studies, review articles and studies performed elsewhere in the world. The remaining references had their full texts (where available) reviewed using a set of strict inclusion criteria, which varied slightly depending on the type of data being reported (i.e. prevalence data or genetic diversity data).

For prevalence data, details on the number of individuals sampled and the different α-thalassaemia alleles and/or genotypes identified were needed. Thus, surveys that reported allele or genotype frequency without the sample size were excluded as were those that reported an overall α-thalassaemia gene frequency without distinguishing between different α-thalassaemia forms or genotypes. Given the important clinical differences between α^+^-, α^0^- and α^ND^-thalassaemia, many of the surveys focused on specific genotypes, in particular those containing an α^0^-thalassaemia allele (i.e. --/αα, --/--, -α/-- and αα^ND^/--). As a result, allele frequencies of the other α-thalassaemia forms were not always a reliable representation of their true allele frequency in the study population, despite them being tested for. In this instance, the reported frequencies of the alleles of secondary interest were entered into the database as missing values. Similarly, if an α-thalassaemia allele was not tested for, the survey was still included but the frequency value of the untested allele was entered as missing. For surveys where the diagnostic algorithm and genotype reporting were sufficiently detailed (e.g. all seven major variants tested using PCR and a range of genotypes reported), we assumed that the lack of explicit reporting of certain genotypes reflected a zero frequency of them. Data on the absence of α-thalassaemia in the studied populations were also included. Where necessary, allele frequencies were derived from their respective genotype frequencies by assuming Hardy-Weinberg proportions.^1, 2^

To be truly representative of the general population, surveys should take place in the community using random sampling. However, it became apparent early in the review process that very few surveys were conducted in this way. We therefore chose to include surveys that sampled from any of the following population groups, provided sampling was random, consecutive or universal: (i) the general community, (ii) pregnant women attending antenatal care and/or their husbands, (iii) cord blood and/or neonates in the absence of any inclusion/exclusion criteria or initial thalassaemia screening, (iv) individuals attending their yearly health check-up, and (v) opt-out volunteers, e.g. students, blood donors or army personnel. These groups were deemed to contain no inherent bias in α-thalassaemia prevalence and thus suitable for the purpose of this study. By contrast, surveys involving opt-in sampling, relatives of known cases of α-thalassaemia, putative anaemic cases, patients or individuals possessing a β-globin variant (i.e. β-thalassaemia, β^E^ or β^S^) were excluded. Moreover, surveys that were carried out in specific ethnic groups were only included if the population being investigated was representative of, or indigenous to, the study area.

Whilst a range of diagnostic methods are available for the diagnosis of α-thalassaemia,^3^ we only included surveys in which molecular diagnostic methods were used (e.g. polymerase chain reaction (PCR)-based techniques and DNA sequencing).^4^ This is because other methods are unreliable for characterising the precise α-thalassaemia genotype of an individual. For instance, haematological indices cannot distinguish between α^+^-thalassaemia and the wild-type state nor between α^0^-thalassaemia and iron deficiency anaemia (both of which lead to microcytosis).^3^ Haemoglobin analysis by capillary electrophoresis (CE), high performance liquid chromatography (HPLC), isoelectric focusing (IEF) and citrate agar electrophoresis (CAE) are all techniques that have been used to identify the quantity of Hb Bart’s, Hb H, Hb CS and other types of haemoglobin relevant to α-thalassaemia, and are comparable in performance.^5, 6^ However, accurate diagnosis of specific α-thalassaemia genotypes is not possible with these techniques and their performance is reduced when there is concomitant inheritance of β-thalassaemia.^5, 7^ Given that the objective of this study was to describe and map allele frequencies, the inclusion of non-molecular studies could affect the reliability of allele frequency estimates.

The majority of surveys that provided information on the genetic diversity of α-thalassaemia mutations were cross-sectional surveys in which there was no prior knowledge or screening of the α-thalassaemia status of the study sample. All of the same inclusion criteria as those described above were applied – that is: (i) a clearly reported sample size and allele or genotype frequencies, (ii) a representative sample, and (iii) molecular diagnosis. In other surveys, the underlying mutations among individuals known to have α-thalassaemia were characterised. To avoid potential bias towards more severe variants in these surveys, only those that took place outside of the hospital setting and in the absence of any inclusion or exclusion criteria pertaining to disease severity were included. Surveys amongst other patient groups, for example β-thalassaemics, were also excluded. Again, only those surveys using molecular diagnostic methods were included.

### Georeferencing

To capture fine-scale spatial heterogeneities in α-thalassaemia gene frequency and/or genetic diversity, included surveys were mapped to the highest spatial resolution possible based on the information provided in the article. Online geopositioning gazeteers were used to identify the latitude and longitude decimal degree coordinates of the study location. For studies that could be georeferenced to the province or district level only, the centroid of the polygon was extracted from ArcMap 10.4.1 (ESRI Inc., Redlands, CA, USA) and used. For studies that were conducted across multiple locations but reported as a single frequency estimate, the coordinates for each site were obtained from the online gazeteers and the centroid of all the sites calculated using ArcMap. In some cases, this resulted in a data point that fell between two land areas, for example Peninsular Malaysia and Malaysian Borneo.

## Supplementary methods 2: Generation of continuous allele frequency maps for Thailand

Continuous maps of the allele frequencies of α^0^-, α^+^- and α^ND^-thalassaemia were generated for Thailand only. This is because there was considerably more data for Thailand than for any of the other countries in the region, in part due to the inclusion of data from local journals. Data from Thailand and all of its neighbouring countries (Myanmar, Lao PDR, Cambodia and Malaysia) were used in this part of the analysis to preclude the possibility of a border effect on the predicted allele frequencies. Surveys that were reported only at the national level were excluded. To generate three separate maps of α-thalassaemia allele frequency, data on each α-thalassaemia form were extracted from the database and used as input for three separate models. The observed allele frequency data were transformed through an empirical logit, which, for databases ≥ 20 surveys, can be well approximated by a Gaussian likelihood.^8^

The goal of the geostatistical analysis was to estimate model parameters and generate predictions of α-thalassaemia allele frequencies in Thailand at fine spatial resolution. We employed, using the “r-inla” package, an easily available Bayesian geostatistical framework involving a stochastic partial differential equation (SPDE) approach^9^ with an integrated nested Laplace approximation (INLA)^10^ algorithm for Bayesian inference. The theoretical principles of this approach have been described in detail elsewhere.^9, 11, 12^ A brief overview is provided below.

Let *Y* be the observed data, which in this study is the set of allele frequency estimates in Thailand and neighbouring countries. Each value *y_i_* represents the allele frequency *y* at location *i*. In general, we can assume that *Y* is generated by an underlying Gaussian process, denoted as *S(x)*, such that any evaluations of *S(x)* are multivariate normal distributed with a given mean and covariance function. Therefore, *S(x)* can be interpreted as a continuously indexed Gaussian field (GF), whereby the effect at each location has a multivariate normal as

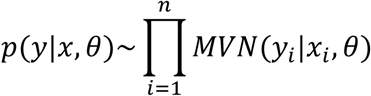

where, *y* denotes allele frequency, *x* is the Gaussian random effect on *y n* represents the number of spatially unique data points, *i* represents location and *θ* denotes the hyperparameters of *x* (i.e. mean *μ* and dispersion parameter *k*). Observations *y* are assumed to be conditionally independent, given *x* and *θ*.

The first law of geography states that: “everything is related to everything else, but near things are more related to distant things.”^13^ This phenomenon is termed spatial dependency, or autocorrelation, and its inclusion in spatial models can greatly improve predictive performance. Several methods for defining the covariance between spatial points exist; here we use the Matérn class of covariance function in which the correlation function is defined based on the Euclidean distance between locations.^14^ Specifically, the stationary Matérn covariance function assumes that if we have two pairs of points separated by the same Euclidean distance *h*, both pairs have same correlation, with correlation monotonically decreasing as a function of *h*.

Matrix algebra operations on a continuously indexed GF are computationally intensive to model. To overcome this, we use a finite element solution of an SPDE approximation to the Matérn covariance function, resulting in a finite dimensional Gaussian Markov random field (GMRF). The finite element representation represents *S(x)* by means of a basis function representation defined on a triangulation of the domain under study, represented as:^9^

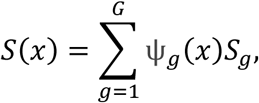

where, *G* is the number of vertices in the triangulation, *ψ_g_*(·) are piecewise polynomial basis functions on each triangle.^15^ The SPDE approach replaces the Matérn covariance function’s dense covariance matrix by a sparse reduced rank matrix. Following the generation of the GMRF prior, the posterior distribution is approximated by an Integrate Nested Laplace Approximation (INLA), which uses a combination of analytical approximation and numerical integration methods to approximate the posterior distribution at each point in the GMRF. The model’s fully Bayesian hierarchical formulation is as follows:

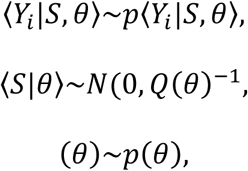

where, again, Y denotes allele frequency (the observation variable), S denotes the underlying Gaussian field, θ denotes a vector of hyperparameters and Q is the sparse precision matrix.

One hundred conditional simulations of the model were performed, whereby all of the pixels in the map were jointly simulated such that spatial autocorrelation between the pixels was accounted for.^16^ This generated PPDs of allele frequency for each pixel in a 1km x 1km grid of Thailand, which were then used to calculate the mean and 95% Bayesian credible intervals for the predicted allele frequencies.

Given the relatively small number of surveys in each dataset, the predictive ability of the model for each form of α-thalassaemia was assessed using a 10-fold cross validation procedure. For each form of α-thalassaemia, the original dataset was randomly partitioned into 10 equal-sized data subsets. In each of 10 separate cross-validation experiments, one of the data subsets was retained as the test set whilst the remaining nine data subsets were used as the training set. The disparity between the model predictions and the observed allele frequencies in the test set was quantified for each cross-validation iteration using: (i) the mean absolute error (MAE), defined as the average magnitude of errors in the predicted values, and (ii) the correlation. The average across the ten cross-validation experiments was then calculated.

R code for the geostatistical model was generated by SB and is available on request. The model was implemented in R using the “r-inla” package, while the predicted allele frequency and disease burden maps were generated in ArcMap.

## Supplementary methods 3: Generating newborn estimates of Hb Bart’s hydrops fetalis and HbH disease for Thailand

Newborn estimates of Hb Bart’s hydrops fetalis and HbH disease (deletional and non-deletional) were generated by pairing our predicted allele frequency PPDs with population and birth rate data. First, the number of births in each pixel of a 1km x 1km grid of Thailand were calculated by multiplying high-resolution 2020 population count data obtained from the WorldPop project website (www.worldpop.org.uk) with the national birth rate. In the UN World Population Prospects 2017 Revision, birth rates were provided as low-, medium- and high-fertility variant projections for 5-year periods. To obtain 2020 estimates of the number of births in Thailand, we used the average of the two 5-year periods 2015-2020 and 2020-2025.

One-hundred realisations of the geostatistical models for α^0^-, α^+^- and α^ND^-thalassaemia allele frequency were run in parallel. For each realisation of the models, the estimated genotype frequencies for Hb Bart’s hydrops fetalis disease (--/--), deletional HbH disease (-α/--) and non-deletional HbH disease (αα^ND^/--) were calculated using the predicted allele frequencies, assuming Hardy-Weinberg proportions for a four-allele system (see Equation 1 in the main text). We also examined the effect that the inclusion of consanguinity into the Hardy-Weinberg equations would have on the estimates (see Table 1 in the main text). Genotype frequencies were then multiplied by the number of births in the corresponding 1km x 1km pixel of the birth count map described above. A PPD for the number of births in each pixel was therefore generated, from which point estimates and uncertainty around these estimates were computed.

The lower bounds of our credible intervals were calculated using the 2.5^th^ percentile for genotype frequency estimates and the low-fertility variant for birth count, whilst the higher bounds were calculated using the 97.5^th^ percentile for genotype frequency estimates and birth count data based on the high-fertility variant.

In order to compare the geostatistical method employed in this study with methods used to generate previous α-thalassaemia disease estimates, we obtained 2003 population count data from the Global Rural-Urban Mapping Project (GRUMP) and paired this with the same birth rate as that used in Modell and Darlison (17 births per 1,000 population).^17^ The same method described above was used; however, no uncertainty intervals were calculated as no corresponding low- and high-fertility variants for birth rate were reported in their study.

## Supplementary methods 4: Generation of maps of genetic variation of α-thalassaemia

From a clinical perspective, the most important distinction between different α-thalassaemia alleles is the number of α-globin genes affected and whether the mutation is of the deletional or non-deletional type – i.e. the distinction between α^0^-, α^+^- and α^ND^-thalassaemia.^18^ However, molecular diagnosis of α-thalassaemia is mutation-specific and as such the profile of α-thalassaemia mutations that are found in a population determines which mutations are tested for in screening programmes. A better understanding of the genetic diversity patterns of α-thalassaemia at the mutation level therefore has important implications for programme design.^4, 19^ In this study, we include surveys that report on genetic diversity at both levels of specificity.

For the descriptive analysis of the relative proportions of each of the three major forms of α-thalassaemia (α^0^-, α^+^ and α^ND^-thalassaemia), only surveys which tested for all three forms were included. This was to avoid misinterpretation of untested forms as being absent. The relative proportions of the different forms were represented using pie charts for which the denominator was the number of α-thalassaemia alleles in the study. The size of the pie charts reflects sample size on the log scale to allow clear visualisation of the data points. A jitter of 0.5-1° latitude and longitude decimal degree coordinates was applied to each survey to avoid the overlapping of surveys carried out in the same location.

The subset of surveys that reported on specific α-thalassaemia mutations indicated that 11 variants were commonly tested for in the Southeast Asia region: single deletion mutations, -α^3.7^ and -α^4.2^; non-deletion mutations, Hb Adana, Hb Constant Spring, Hb Paksé and Hb Quong Sze; and double deletion mutations, (α)20.5, FIL, MED, SEA and THAI. Other non-deletion mutations such as Hb Q-Thailand and initiation codon mutations were tested for in very few surveys (*n* = 13) and were grouped together into a single category as “Other”. Surveys were mapped spatially using bar charts to depict variant proportions. All variants that were tested for in a survey were represented along the *x*-axis of the corresponding chart. Empty spaces are therefore indicative of the variant being absent in the sample. To allow clear visualisation of the bar charts, data for Thailand were displayed separately, whilst the remaining Southeast Asian countries were divided into two maps: (i) Myanmar, Lao PDR, Cambodia and Vietnam in the north, and (ii) Malaysia, Indonesia and Singapore in the south. Bar charts could not be placed at their precise geographical location and so surveys were mapped as points and connected to their corresponding bar chart by a black line.

## Supplementary results: Existing α-thalassaemia maps and characteristics of α-thalassaemia database

Among the surveys included in the final geodatabase, α^0^-thalassaemia was the most extensively studied form (*n* = 97), followed by α^ND^-thalassaemia (*n* = 50) and then α^+^-thalassaemia (*n* = 47). The majority of studies (95.3%) were carried out after 1995 and in particular from 2005 onwards (78.3%) (Supplementary figure 2B). In terms of the population groups sampled, 21.7% of surveys were community-based, 59.4% were carried out amongst selected but unbiased population groups (e.g. pregnant women, husbands of pregnant women, neonates and cord blood) and 8.5% in opt-out volunteers. One (<1%) survey providing only genetic diversity data was carried out in individuals known to have α-thalassaemia. Twelve per cent of surveys provided no detailed information on the sampling methodology but were judged to be unbiased (based on other information provided in the article or personal communication with the corresponding author) and were included. Sample size ranged from 4 to 55,796. Small sample sizes (e.g. *n* = 4) came from surveys that were carried out across multiple geographic locations and the α-thalassaemia frequencies reported separately for each. Supplementary table 1 provides a breakdown of the survey features for the overall database and for prevalence surveys and genetic variant surveys, separately.

Supplementary figure 2 here.

Supplementary figure 3 here.

Supplementary table 1 here.

Supplementary table 2 here.

## Supplementary figures and tables

**Supplementary figure 1** A map of the countries included in this study. Here we defined the Southeast Asian region according to the member states of the Association of Southeast Asian Nations (ASEAN) (*http://asean.org/asean/asean-member-states/*).

**Supplementary figure 2** Spatial and temporal distributions of the α-thalassaemia surveys included in the final database. In both panels, the shape of the data points indicates the type of data provided by the survey, the colour indicates whether the survey was found in our online literature search or in local journals, and size represents the sample size of the survey. In (A) a spatial jitter of up to 0.3^0^ latitude and longitude decimal degree coordinates was applied to allow visualisation of spatially duplicated data points.

**Supplementary figure 3** A map of our current knowledge of the global distribution, gene frequency and genetic diversity of α-thalassemia. Only the most common variants for α^+^-thalassemia (-α^3.7^and -α^4.2^) and α^0^-thalassemia (--^MED^ and --^SEA^) are shown for each region. The variants that appear in parentheses are those for which the data used to make this map are limited. Figure taken from Piel and Weatherall (2014).^18^

**Supplementary table 1** Summary of the α-thalassaemia dataset characteristics according to the type of data provided (allele frequency data or genetic variant data), and overall. Numbers correspond to individual surveys that met the study inclusion criteria. As some sources reported more than one survey from multiple locations or in multiple population groups, the number of surveys is greater than the number of references in the Supplementary bibliographies S1 and S2. Some surveys reported data on both α-thalassaemia prevalence and genetic diversity and are therefore included twice in these columns, but once in the overall column.

**Supplementary table 2** Observed allele frequency ranges for different α-thalassaemia forms.

Sources from which the data points used in the allele maps were identified included:

Sources from which the data points used in the genetic variation maps were identified included:

## References

1. Piel FB, Weatherall DJ. The alpha-thalassemias. The New England journal of medicine. 2014;371(20):1908–16.

2. Flint J, Harding RM, Boyce AJ, Clegg JB. The population genetics of the haemoglobinopathies. Bailliere’s clinical haematology. 1998;11(1):1–51.

3. Premawardhena A, Allen A, Piel F, Fisher C, Perera L, Rodrigo R, et al. The evolutionary and clinical implications of the uneven distribution of the frequency of the inherited haemoglobin variants over short geographical distances. British journal of haematology. 2017;176(3):475–84.

4. Weatherall DJ, Clegg JB. Distribution and Population Genetics of the Thalassaemias. The Thalassaemia Syndromes: Blackwell Science Ltd; 2001. p. 237–84.

5. May J, Evans JA, Timmann C, Ehmen C, Busch W, Thye T, et al. Hemoglobin variants and disease manifestations in severe falciparum malaria. JAMA. 2007;297(20):2220–6.

6. Vichinsky EP. Clinical Manifestations of a-Thalassemia. Cold Spring Harbor Perspectives in Medicine. 2013;3(5):a011742.

7. Hoppe CC. Newborn screening for non-sickling hemoglobinopathies. Hematology American Society of Hematology Education Program. 2009:19–25.

8. Weatherall DJ, Clegg JB. The Pathophysiology of the Thalassaemias. The Thalassaemia Syndromes: Blackwell Science Ltd; 2001. p. 192–236.

9. Galanello R, Cao A. Alpha-thalassemia.

10. Chui DH, Fucharoen S, Chan V. Hemoglobin H disease: not necessarily a benign disorder. Blood. 2003;101(3):791–800.

11. Fucharoen S, Viprakasit V. Hb H disease: clinical course and disease modifiers. Hematology American Society of Hematology Education Program. 2009:26–34.

12. Lal A, Goldrich ML, Haines DA, Azimi M, Singer ST, Vichinsky EP. Heterogeneity of Hemoglobin H Disease in Childhood. New England Journal of Medicine. 2011;364(8):710–8.

13. Weatherall DJ, Wood WG, Pressley L, Higgs DR, Clegg JB. Molecular Basis for Mild Forms of Homozygous Beta-Thalassaemia. The Lancet. 1981;317(8219):527–9.

14. Wainscoat JS, Kanavakis E, Wood WG, Letsky EA, Huehns ER, Marsh GW, et al. Thalassaemia intermedia in Cyprus: the interaction of alpha and beta thalassaemia. British journal of haematology. 1983;53(3):411–6.

15. Wai Kan Y, G. Nathan D. Mild thalassemia: the result of interactions of alpha and beta thalassemia genes 1970. 635–42 p.

16. Law HY, Chee MK, Tan GP, Ng IS. The simultaneous presence of alpha- and beta-thalassaemia alleles: a pitfall of thalassaemia screening. Community genetics. 2003;6(1):14–21.

17. Weatherall DJ, Akinyanju O, Fucharoen S, Olivieri N, Musgrove P. Inherited Disorders of Hemoglobin. Disease Control Priorities in Developing Countries. 2nd edition ed. New York: Oxford University Press; 2006.

18. Chui DHK. Alpha-thalassaemia and population health in Southeast Asia. Annals of Human Biology. 2005;32(2):123–30.

19. Songdej D, Babbs C, Higgs DR. An international registry of survivors with Hb Bart’s hydrops fetalis syndrome. Blood. 2017;129(10):1251–9.

20. Fucharoen S, Winichagoon P. Hemoglobinopathies in Southeast Asia. Hemoglobin. 1987;11(1):65–88.

21. Fucharoen S, Winichagoon P. Haemoglobinopathies in southeast Asia. Indian J Med Res. 2011;134:498–506.

22. Modell B, Darlinson M. Global epidemiology of haemoglobin disorders and derived service indicators. Bulletin of the World Health Organization. 2008;86(6):480–7.

23. Piel FB, Howes RE, Patil AP, Nyangiri OA, Gething PW, Bhatt S, et al. The distribution of haemoglobin C and its prevalence in newborns in Africa. Scientific Reports. 2013;3:1671.

24. Piel FB, Patil AP, Howes RE, Nyangiri OA, Gething PW, Dewi M, et al. Global epidemiology of sickle haemoglobin in neonates: a contemporary geostatistical model-based map and population estimates. Lancet. 2013;381(9861):142–51.

25. Howes RE, Dewi M, Piel FB, Monteiro WM, Battle KE, Messina JP, et al. Spatial distribution of G6PD deficiency variants across malaria-endemic regions. Malar J. 2013;12:418.

26. Munkongdee T, Tanakulmas J, Butthep P, Winichagoon P, Main B, Yiannakis M, et al. Molecular Epidemiology of Hemoglobinopathies in Cambodia. Hemoglobin. 2016;40(3):163–7.

27. Sengchanh S, Sanguansermsri T, Horst D, Horst J, Flatz G. High frequency of alpha-thalassemia in the So ethnic group of south Laos. Acta Haematologica. 2005;114(3):164– 6.

28. Alauddin H, Langa M, Mohd Yusoff M, Raja Sabudin RZA, Ithnin A, Abdul Razak NF, et al. Detection of alpha-thalassaemia in neonates on cord blood and dried blood spot samples by capillary electrophoresis. The Malaysian journal of pathology. 2017;39(1):17– 23.

29. Tan Jama, Lee PC, Wee YC, Tan KL, Mahali NF, George E, et al. High prevalence of alpha- and beta-thalassemia in the kadazandusuns in east Malaysia: Challenges in providing effective health care for an indigenous group. Journal of Biomedicine and Biotechnology. 2010;2010.

30. Weatherall D. The inherited disorders of haemoglobin: an increasingly neglected global health burden. The Indian Journal of Medical Research. 2011;134(4):493–7.

31. Ibrahim H. Current situation in control strategies and health systems in Asia - Malaysia. Malaysia: Ministry of Health 2012.

32. Blencowe H, Cousens S, Jassir FB, Say L, Chou D, Mathers C, et al. National, regional, and worldwide estimates of stillbirth rates in 2015, with trends from 2000: a systematic analysis. The Lancet Global Health.4(2):e98–e108.

33. World population prospects: The 2017 revision. [Internet]. [cited July 27 2017]. Available from: https://esa.un.org/unpd/wpp/.

34. Liebhaber SA, Kan YW. Differentiation of the mRNA transcripts originating from the alpha 1- and alpha 2-globin loci in normals and alpha-thalassemics. Journal of Clinical Investigation. 1981;68(2):439–46.

35. Orkin SH, Goff SC, Hechtman RL. Mutation in an intervening sequence splice junction in man. Proc Natl Acad Sci U S A. 1981;78(8):5041–5.

36. Kulaphisit M, Kampuansai J, Leecharoenkiat K, Wathikthinnakon M, Kangwanpong D, Munkongdee T, et al. A comprehensive ethnic-based analysis of alpha thalassaemia allelle frequency in northern Thailand. Sci Rep. 2017;7(1):4690.

37. Douglas NM, Anstey NM, Buffet PA, Poespoprodjo JR, Yeo TW, White NJ, et al. The anaemia of Plasmodium vivax malaria. Malaria Journal. 2012;11(1):135.

38. Old J, Henderson S. Molecular diagnostics for haemoglobinopathies. Expert Opinion on Medical Diagnostics. 2010;4(3):225–40.

39. Ratanasiri T, Komwilaisak R, Sittivech A, Kleebkeaw P, Seejorn K. Incidence, causes and pregnancy outcomes of hydrops fetalis at Srinagarind Hospital, 1996-2005: a 10-year review. Journal of the Medical Association of Thailand = Chotmaihet thangphaet. 2009;92(5):594–9.

40. Galanello R. Screening and Diagnosis for Haemoglobin Disorders. In: Old J, editor. Prevention of Thalassaemias and Other Hemoglobin Disorders. 1. Nicosia, Cyprus: Thalassaemia International Federation; 2013.

41. Wong D. The modifiable areal unit problem (MAUP). The SAGE handbook of spatial analysis. 2009:105–23.

42. Fucharoen S, Weatherall DJ. The hemoglobin E thalassemias. Cold Spring Harb Perspect Med. 2012;2(8).

43. Viprakasit V, Tanphaichitr VS, Chinchang W, Sangkla P, Weiss MJ, Higgs DR. Evaluation of alpha hemoglobin stabilizing protein (AHSP) as a genetic modifier in patients with beta thalassemia. Blood. 2004;103(9):3296–9.

44. Bundhamcharoen K, Odton P, Phulkerd S, Tangcharoensathien V. Burden of disease in Thailand: changes in health gap between 1999 and 2004. BMC Public Health. 2011;11(1):53.

45. Dhillon PK, Jeemon P, Arora NK, Mathur P, Maskey M, Sukirna RD, et al. Status of epidemiology in the WHO South-East Asia region: burden of disease, determinants of health and epidemiological research, workforce and training capacity. International journal of epidemiology. 2012;41(3):847–60.

46. Rue H, Martino S, Chopin N. Approximate Bayesian inference for latent Gaussian models by using integrated nested Laplace approximations. Journal of the Royal Statistical Society: Series B (Statistical Methodology). 2009;71(2):319–92.

47. Hardy GH. Mendelian Proportions in a Mixed Population. Science (New York, NY). 1908;28(706):49–50.

48. Weinberg W. Über den Nachweis der Vererbung beim Menschen. Jahresh Wuertt Verh Vaterl Naturkd1908. p. 369–82.

49. Tatem AJ. WorldPop, open data for spatial demography. 2017;4:170004.

50. Liu YT, Old JM, Miles K, Fisher CA, Weatherall DJ, Clegg JB. Rapid detection of alpha-thalassaemia deletions and alpha-globin gene triplication by multiplex polymerase chain reactions. British journal of haematology. 2000;108(2):295–9.

## Supplementary bibliography S1

1. Hardy GH. Mendelian Proportions in a Mixed Population. Science (New York, NY) 1908; 28(706): 49–50.

2. Weinberg W. Über den Nachweis der Vererbung beim Menschen. Jahresh Wuertt Verh Vaterl Naturkd; 1908: 369–82.

3. Fucharoen S, Winichagoon P. Haemoglobinopathies in southeast Asia. Indian J Med Res 2011; 134: 498–506.

4. Old J, Henderson S. Molecular diagnostics for haemoglobinopathies. Expert Opinion on Medical Diagnostics 2010; 4(3): 225–40.

5. Alauddin H, Langa M, Mohd Yusoff M, et al. Detection of alpha-thalassaemia in neonates on cord blood and dried blood spot samples by capillary electrophoresis. The Malaysian journal of pathology 2017; 39(1): 17–23.

6. Mantikou E, Arkesteijn Sg Fau - Beckhoven van JM, Beckhoven van Jm Fau - Kerkhoffs J-L, Kerkhoffs Jl Fau - Harteveld CL, Harteveld Cl Fau - Giordano PC, Giordano PC. A brief review on newborn screening methods for hemoglobinopathies and preliminary results selecting beta thalassemia carriers at birth by quantitative estimation of the HbA fraction. (1873-2933 (Electronic)).

7. Chui DHK. Alpha-thalassaemia and population health in Southeast Asia. Annals of Human Biology 2005; 32(2): 123–30.

8. Diggle PJ, Ribeiro PJ. Model-based Geostatistics. 1 ed: Springer-Verlag New York; 2007.

9. Lindgren F, Rue H, Lindström J. An explicit link between Gaussian fields and Gaussian Markov random fields: the stochastic partial differential equation approach. Journal of the Royal Statistical Society: Series B (Statistical Methodology) 2011; 73(4): 423–98.

10. Rue H, Martino S, Chopin N. Approximate Bayesian inference for latent Gaussian models by using integrated nested Laplace approximations. Journal of the Royal Statistical Society: Series B (Statistical Methodology) 2009; 71(2): 319–92.

11. Rue H, Riebler A, Sørbye SH, Illian J, Simpson DP, Lindgren F. Bayesian Computing with INLA: A Review. Annual Review of Statistics and its Application 2017; 4(1).

12. Cameletti M, Lindgren F, Simpson D, Rue H. Spatio-temporal modeling of particulate matter concentration through the SPDE approach. AStA Advances in Statistical Analysis 2013; 97(2): 109–31.

13. Tobler WR. A Computer Movie Simulating Urban Growth in the Detroit Region. Economic Geography 1970; 46: 234–40.

14. Krainski ET, Lindgren F, Simpson D, Rue R. The R-INLA tutorial on SPDE models. 2017. http://www.math.ntnu.no/inla/r-inla.org/tutorials/spde/R/spde-tutorial-introduction.R.

15. Moraga P, Cano J, Baggaley RF, et al. Modelling the distribution and transmission intensity of lymphatic filariasis in sub-Saharan Africa prior to scaling up interventions: integrated use of geostatistical and mathematical modelling. Parasites & Vectors 2015; 8(1): 560.

16. Gething PW, Patil AP, Hay SI. Quantifying Aggregated Uncertainty in Plasmodium falciparum Malaria Prevalence and Populations at Risk via Efficient Space-Time Geostatistical Joint Simulation. PLOS Computational Biology 2010; 6(4): e1000724.

17. Modell B, Darlinson M. Global epidemiology of haemoglobin disorders and derived service indicators. Bulletin of the World Health Organization 2008; 86(6): 480–7.

18. Piel FB, Weatherall DJ. The alpha-thalassemias. The New England journal of medicine 2014; 371(20): 1908–16.

19. Galanello R. Screening and Diagnosis for Haemoglobin Disorders. In: Old J, ed. Prevention of Thalassaemias and Other Hemoglobin Disorders. Nicosia, Cyprus: Thalassaemia International Federation; 2013.

## Supplementary bibliography S2: Allele frequency data

1. Alauddin H, Langa M, Mohd Yusoff M, et al. Detection of alpha-thalassaemia in neonates on cord blood and dried blood spot samples by capillary electrophoresis. The Malaysian journal of pathology 2017; 39(1): 17–23.

2. Amornkitbamrung S. Screening for Thalassemia Disease, α-thalassemia 1 trait, ß-thalassemia trait and Hb E in pregnant women at Nongkhai Hospital. Chonburi Hospital Journal 2005; 30(3): 171–86.

3. Apidechkul T. Prevalence of thalassemia carriers among the Lahu hill tribe population, Chiang Rai, Thailand. Asian Biomedicine 2015; 9(4): 527–33.

4. Ausavarungnirun R, Winichagoon P, Fucharoen S, Epstein N, Simkins R. Detection of zeta-globin chains in the cord blood by ELISA (enzyme-linked immunosorbent assay): Rapid screening for alpha-thalassemia 1 (Southeast Asian type). Am J Hematol 1998; 57(4): 283–6.

5. Chareonkul P, Kraisin J. Prevention and control of thalassemia at Saraburi Regional Hospital. J Med Assoc Thai 2004; 1(8): 8–15.

6. Chinorose S, Toongkam K. Prevalence of Thalassemai Trait and Identification of Couples Risk in Pregnant Women for Severe Thalassemia Disease in Phayao Hospital. Uttaradit Hospital Medical Bulletin 2009; 24(1): 61–9.

7. Dangwibul S, al. e. Efficiency of severe thalassemia screeing program in 5 rural hospitals in Roi Et province (Oral presentation, abstract). The 8th National Thalassemia Academic Symposium. Khon Kaen; 2002. p. 174-5.

8. Fucharoen G, al. e. Thalassemia and iron deficiency in subjects with positive screening OF test and KKU-DCIP-Clear. Thai J Hematol Transf Med 1999; 9: 111–8.

9. Fucharoen G, Sanchaisuriya K, Sae-ung N, Dangwibul S, Fucharoen S. A simplified screening strategy for thalassaemia and haemoglobin E in rural communities in south-east Asia. Bulletin of the World Health Organization 2004; 82(5): 364–72.

10. Fucharoen S, Winichagoon P, Wisedpanichkij R, et al. Prenatal and postnatal diagnoses of thalassemias and hemoglobinopathies by HPLC. Clin Chem 1998; 44(4): 740–8.

11. Hundrieser J, Laig M, Yongvanit P, et al. Study of Alpha-Thalassemia in Northeastern Thailand at the DNA Level. Hum Hered 1990; 40(2): 85–8.

12. Hundrieser J, Sanguansermsri T, Papp T, Flatz G. Alpha-Thalassemia in Northern Thailand - Frequency of Deletional Types Characterized at the DNA Level. Hum Hered 1988; 38(4): 211–5.

13. Jameela S, Sabirah SO, Babam J, et al. Thalassaemia screening among students in a secondary school in Ampang, Malaysia. The Medical journal of Malaysia 2011; 66(5): 522–4.

14. Jearakul W, Khamsaen J. Alpha-thalassemia 1 among Married Couples in Six Northeastern Provinces. Journal of Health Science 2009; 18(5): 728–35.

15. Jindatanmanusan P, Riolueang S, Glomglao W, et al. Diagnostic applications of newborn screening for alpha-thalassaemias, haemoglobins E and H disorders using isoelectric focusing on dry blood spots. Annals of clinical biochemistry 2013.

16. Jopang Y. Thalassemia and hemoglobinopathies in anemic schoolchildren The 13th National Thalassemia Academic Symposium. Bangkok; 2007. p. Page 154.

17. Jopang Y. Thalassemia and hemoglobinopathies in anemic schoolchildren. Health Promoting Hospital, Regional Health Promotion Center 5. Nakhonratchasima. Journal of the Medical Technologist Association of Thailand 2008; 36(1): 2235–41.

18. Jopang Y, Mernkratok S, Puangpiruk R. The efficacy of thalassemias and Hb E screening in first-trimester pregnant women at Health Promoting Hospital, Regional Health Promotion Center 5 Nakhonratchasima (poster abstract). The 10th National Thalassemia Academic Symposium. Bangkok; 2004. p. 171.

19. Karakochuk CD, Whitfield KC, Barr SI, et al. Genetic hemoglobin disorders rather than iron deficiency are a major predictor of hemoglobin concentration in women of reproductive age in rural prey veng, Cambodia. Journal of Nutrition 2015; 145(1): 134–42.

20. Karnpean R, Pansuwan A, Fucharoen G, Fucharoen S. Evaluation of the URIT-2900 Automated Hematology Analyzer for screening of thalassemia and hemoglobinopathies in Southeast Asian populations. Clin Biochem 2011; 44(10-11): 889–93.

21. Koh DXR, Raja Sabudin RZA, Mohd Yusoff M, et al. Molecular Characterisation of alpha- and beta-Thalassaemia among Indigenous Senoi Orang Asli Communities in Peninsular Malaysia. Annals of human genetics 2017.

22. LemmensZygulska M, Eigel A, Helbig B, Sanguansermsri T, Horst J, Flatz G. Prevalence of alpha-thalassemias in northern Thailand. Hum Genet 1996; 98(3): 345–7.

23. Limsakulsiriratt P, Oncoung W. High Risk Couples for Hb Bart’s Hydrops Fetalis in Public Health Region 8 and 9 During 2004-2006. Chonburi Hospital Journal 2007; 32(1): 9–14.

24. Munkongdee T, Pichanun D, Butthep P, et al. Quantitative analysis of Hb Bart’s in cord blood by capillary electrophoresis system. Ann Hematol 2011; 90(7): 741–6.

25. Munkongdee T, Tanakulmas J, Butthep P, et al. Molecular Epidemiology of Hemoglobinopathies in Cambodia. Hemoglobin 2016; 40(3): 163–7.

26. Nguyen HV, Sanchaisuriya K, Nguyen D, et al. Thalassemia and Hemoglobinopathies in Thua Thien Hue Province, Central Vietnam. Hemoglobin 2013; 37(4): 333–42.

27. Nguyen NT, Sanchaisuriya K, Sanchaisuriya P, et al. Thalassemia and hemoglobinopathies in an ethnic minority group in Central Vietnam: implications to health burden and relationship between two ethnic minority groups. Journal of Community Genetics 2017: 1– 8.

28. Nguyen VH, Sanchaisuriya K, Wongprachum K, et al. Hemoglobin Constant Spring is markedly high in women of an ethnic minority group in Vietnam: a community-based survey and hematologic features. Blood cells, molecules & diseases 2014; 52(4): 161–5.

29. Nillakupt K, Nathalang O, Arnutti P, Jindadamrongwech SB T., Panichkul S, Areekul W. Prevalence and hematological parameters of thalassemia in Tha Kradarn subdistrict Chachoengsao Province, Thailand. J Med Assoc Thai 2012; 95(Suppl 5): S124–S32.

30. O’Riordan S, Hien TT, Miles K, et al. Large scale screening for haemoglobin disorders in southern Vietnam: implications for avoidance and management. British journal of haematology 2010; 150(3): 359–64.

31. Panomai N, Sanchaisuriya K, Yamsri S, et al. Thalassemia and iron deficiency in a group of northeast Thai school children: relationship to the occurrence of anemia. Eur J Pediatr 2010; 169(11): 1317–22.

32. Panyasai S, Cheechang S. The efficiency of screening for carriers of severe thalassemia in three community hospitals in Nakhon Si Thammarat province, Thailand. Songkla Med J 2009; 27(1): 61–72.

33. Panyasai S, Sringam P, Fucharoen G, Sanchaisuriya K, Fucharoen S. A simplified screening for alpha-thalassemia 1 (SEA type) using a combination of a modified osmotic fragility test and a direct PCR on whole blood cell lysates. Acta Haematol-Basel 2002; 108(2): 74–8.

34. Pharephan S, Sirivatanapa P, Makonkawkeyoon S, Tuntiwechapikul W, Makonkawkeyoon L. Prevalence of α-thalassaemia genotypes in pregnant women in northern Thailand. Indian Journal of Medical Research 2016; 143(MARCH): 315–22.

35. Phollarp P, Tritipsombut J, Worasan C, et al. Thalassemia and iron deficiency among pregnant women attending antenatal care service at Khao Wong Hospital, Kalasin province. Journal of Medical Technology and Physical Therapy 2010; 22(3): 262–70.

36. Pichanun D, Munkongdee T, Klamchuen S, et al. Molecular Screening of the Hbs Constant Spring (codon 142, TAA > CAA, alpha 2) and Pakse (codon 142, TAA > TAT, alpha 2) Mutations in Thailand. Hemoglobin 2010; 34(6): 582–6.

37. Rahimah AN, Nisha S, Safiah B, et al. Distribution of alpha thalassaemia in 16 year old Malaysian Students in Penang, Melaka and Sabah. The Medical journal of Malaysia 2012; 67(6): 565–70.

38. Rasri W. Prevention and control of seven thalassemia in pregnancy at Phayao Hospital. Journal of Health Science 2008; 17: 477–84.

39. Rawangkran A, Janwithee N, Wong P, Jermnim N. Prevalence of Thalassemia Trait from Screening Program in Pregnant Women in the Lower Northern Region of Thailand. Thai J Genet 2013; 1: 156–9.

40. Sanchaisuriya K, Fucharoen S, Ratanasiri T, et al. Thalassemia and hemoglobinopathies rather than iron deficiency are major causes of pregnancy-related anemia in northeast Thailand. Blood Cell Mol Dis 2006; 37(1): 8–11.

41. Sangnark P. Prevalence of thalassemia and hemoglobinopathies in pregnant women at Bangkrathum Hospital, Phitsanulok province. Buddhachinaraj Med J 2009; 26(1): 36–43.

42. Sanguansermsri T, Phumyu N, Chomchuen S, Steger HF. Screening for alpha-thalassemia-1 heterozygotes in expecting couples by the combination of a simple erythrocyte osmotic fragility test and a PCR-based method. Community Genetics 1999; 2(1): 26–9.

43. Sanguansermsri T, Steger HF, Sirivatanapa P, Wanapirak C, Tongsong T. Prevention and Control of Severe Thalassemia Syndrome: Chiang Mai strategy. Thai J Hematol Transf Med 1998; 8: 207–14.

44. Sattaratanamai C, Thongsutti S, Sucharitcheep P, Tuengsaeng D, Chomchuen S. Prevalence of Thalassemia and Hemoglobinopathies in Pregnant women at Surin Hospital. Medical Journal of Srisaket Surin Buriram Hospitals 2000; 15(1): 1–12.

45. Savongsy O, Fucharoen S, Fucharoen G, Sanchaisuriya K, Sae-ung N. Thalassemia and hemoglobinopathies in pregnant Lao women: carrier screening, prevalence and molecular basis. Ann Hematol 2008; 87(8): 647–54.

46. Sengchanh S, Sanguansermsri T, Horst D, Horst J, Flatz G. High frequency of alpha-thalassemia in the So ethnic group of south Laos. Acta Haematol-Basel 2005; 114(3): 164– 6.

47. Sirichotiyakul S, Tantipalakorn C, Sanguansermsri T, Wanapirak C, Tongsong T. Erythrocyte osmotic fragility test for screening of alpha-thalassemia-1 and beta-thalassemia trait in pregnancy. Int J Gynecol Obstet 2004; 86(3): 347–50.

48. Soonklang M, Nonthalee S, Juntharaniyom M. Thalassemia and hemoglobinopathies in couples, Khon Kaen Hospital (Poster abstract). The 20th National Thalassemia Academic Symposium. Bangkok; 2014. p. Page P29.

49. Sornkayasit K, al. e. Incidence of Hb Constant Spring and Hb Pakse in Khon Kaen: Using capillary electrophoresis and DNA analysis (Poster abstract). The 18th National Thalassemia Academic Symposium. Nonthaburi; 2012. p. 90.

50. Srivorakun H, Fucharoen G, Changtrakul Y, Komwilaisak P, Fucharoen S. Thalassemia and hemoglobinopathies in Southeast Asian newborns: diagnostic assessment using capillary electrophoresis system. Clin Biochem 2011; 44(5-6): 406–11.

51. Sukrat B, Sirichotiyakul S. The prevalence and causes of anemia during pregnancy in Maharaj Nakorn Chiang Mai Hospital. J Med Assoc Thai 2006 oct;89 Suppl 4:S142–6 2006; **89**(Suppl 4): S142-6.

52. Sutjasung P, Fucharoen G, Fucharoen S, Chattumaruk P, Changtrakun D, Sanchaisuriya K. Effectiveness of thalassemia screening with the use of internal quality control blood samples at Kasetsomboon Hospital, Chaiyaphoom province. Journal of Medical Technology and Physical Therapy 2011; 23(1): 34–45.

53. Suwannakhon N, Seeratanachot T, Mahingsa K, Namwong P, T. S. Prevalence of Alpha-thalassemia Trait in the Volunteered Personals of University of Phayao. J Hematol Transfus Med 2014; 24: 129–36.

54. Tan JA, Tay JS, Soemantri A, et al. Deletional types of alpha-thalassaemia in central Java. Hum Hered 1992; 42(5): 289–92.

55. Tan Jama, Lee PC, Wee YC, et al. High prevalence of alpha- and beta-thalassemia in the kadazandusuns in east Malaysia: Challenges in providing effective health care for an indigenous group. Journal of Biomedicine and Biotechnology 2010; 2010.

56. Tangvarasittichai O, Jeenapongsa R, Sitthiworanan C, Sanguansermsri T. Laboratory investigations of Hb Constant Spring. Clin Lab Haematol 2005; 27(1): 47–9.

57. Tangvarasittichai O, Poonanan N, Tangvarasittichai S. Using Red Cell Indices and Reticulocyte Parameters for Carrier Screening of Various Thalassemia Syndromes. Indian journal of clinical biochemistry: IJCB 2017; 32(1): 61–7.

58. Tanphaichitr VS, Pung-amritt P, Puchaiwatananon O, et al. Studies on hemoglobin Bart’s and deletion of alpha-globin genes from cord blood in Thailand (poster abstract). The International Conference on Thalassemia. Bangkok; 1985. p. P05.

59. Than AM, Harano T, Harano K, Myint AA, Ogino T, Okada S. High incidence of alpha-thalassemia, hemoglobin E, and glucose-6-phosphate dehydrogenase deficiency in populations of malaria-endemic southern Shan State, Myanmar. Int J Hematol 2005; 82(2): 119–23.

60. Thanomrat P, Wannapira W, Sritippawan S, Boon-eam O. The prevalence of alpha-thalassemia trait in Buddhachinaraj Phitsanulok Hospital preliminary report. Buddhachinaraj Med J 2003; 20(1): 19–25.

61. Thongon R, al. e. Strategy for thalassemia screening at Yala hospital (Poster abstract). The 15 th National Thalassemia Academic Symposium. Udon Thani; 2009. p. 144.

62. Tienthavorn V, Pattanapongsthorn J, Charoensak S, Sae-Tung R, Charoenkwan P, Sanguansermsri T. Prevalence of Thalassemia Carriers in Thailand. Thai J Hematol Transf Med 2006; 16: 307–12.

63. Tongon R, Yunu R, Sanchaisuriya K, et al. Thalassemia and hemoglobinopathies in pregnant women attended antenatal care service at Yala Hospital. J Med Tech Phy Ther 2014; 26(1): 32–9.

64. Traisrisilp K, Jatavan P, Tongsong T. A retrospective comparison of pregnancy outcomes between women with alpha-thalassaemia 1 trait and normal controls. Journal of obstetrics and gynaecology: the journal of the Institute of Obstetrics and Gynaecology 2017: 1–4.

65. Trisakul N, Siripulsak P, Chuesupalobol W. Prevalence of Thalassemia, Hemoglobinopathy, At-Risk Couples and Incidence of Thalassemia Major from the Screening Program, Prenatal and Postnatal Diagnosis at Yasothorn Hospital. J Hematol Transfus Med 2008; 19: 285–92.

66. Tritipsombut J, Sanchaisuriya K, Fucharoen S, et al. Hemoglobin Profiles and Hematologic Features of Thalassemic Newborns Application to Screening of alpha-Thalassemia 1 and Hemoglobin E. Arch Pathol Lab Med 2008; 132(11): 1739–45.

67. Tritipsombut J, Sanchaisuriya K, Phollarp P, et al. Micromapping of Thalassemia and Hemoglobinopathies in Diferent Regions of Northeast Thailand and Vientaine, Laos People’s Democratic Republic. Hemoglobin 2012; 36(1): 47–56.

68. Uaprasert N, Settapiboon R, Amornsiriwat S, et al. Diagnostic utility of isoelectric focusing and high performance liquid chromatography in neonatal cord blood screening for thalassemia and non-sickling hemoglobinopathies. Clinica chimica acta; international journal of clinical chemistry 2014; 427: 23–6.

69. Wanapirak C, Muninthorn W, Sanguansermsri T, Dhananjayanonda P, Tongsong T. Prevalence of Thalassemia in pregnant women at Maharaj Nakorn Chiang Mai Hospital. Journal of the Medical Association of Thailand 2004; 87(12): 1415–8.

70. Wong P, Thanormrat P, Srithipayawan S, et al. Risk of a couple having a child with severe thalassemia syndrome, prevalence in lower northern Thailand. Southeast Asian Journal of Tropical Medicine and Public Health 2006; 37(2): 366–9.

71. Wong P, Thanormrat P, Srithipayawan S, et al. Prevalence of thalassemia trait from screening program in pregnant women of Phitsanulok. Thai J Hematol Transf Med 2004; 14: 181–6.

72. Wongkham J, Ratanasiri T, Komwilaisak R, Saksiriwuttho P, Paibool M, Chatvised P. Thalassemia screening in pregnant women at antenatal care clinic, Srinagarind Hospital. Srinagarind Med J 2013; 28(2): 170–7.

73. Yap ZM, Sun KM, Teo CRL, Tan ASC, Chong SS. Evidence of differential selection for the -alpha(3.7) and -alpha(4.2) single-alpha-globin gene deletions within the same population. Eur J Haematol 2013; 90(3): 210–3.

74. Yin SKK, Chong QT, Mei LA, et al. A molecular epidemiologic study of thalassemia using newborns’ cord blood in a multiracial Asian population in Singapore - Results and recommendations for a population screening program. J Pediat Hematol Onc 2004; 26(12): 817–9.

## Supplementary bibliography S3: Genetic variant data

2. Fucharoen G, al. e. Thalassemia and iron deficiency in subjects with positive screening of test and KKU-DCIP-Clear. Thai J Hematol Transf Med 1999; 9: 111–8.

3. Fucharoen G, Sanchaisuriya K, Sae-ung N, Dangwibul S, Fucharoen S. A simplified screening strategy for thalassaemia and haemoglobin E in rural communities in south-east Asia. Bulletin of the World Health Organization 2004; 82(5): 364–72.

4. Fucharoen S, Winichagoon P, Wisedpanichkij R, et al. Prenatal and postnatal diagnoses of thalassemias and hemoglobinopathies by HPLC. Clin Chem 1998; 44(4): 740–8.

5. Hundrieser J, Laig M, Yongvanit P, et al. Study of Alpha-Thalassemia in Northeastern Thailand at the DNA Level. Hum Hered 1990; 40(2): 85–8.

6. Hundrieser J, Sanguansermsri T, Papp T, Flatz G. Alpha-Thalassemia in Northern Thailand - Frequency of Deletional Types Characterized at the DNA Level. Hum Hered 1988; 38(4): 211–5.

7. Jameela S, Sabirah SO, Babam J, et al. Thalassaemia screening among students in a secondary school in Ampang, Malaysia. The Medical journal of Malaysia 2011; 66(5): 522–4.

8. Jearakul W, Khamsaen J. Alpha-thalassemia 1 among Married Couples in Six Northeastern Provinces. Journal of Health Science 2009; 18(5): 728–35.

9. Jindatanmanusan P, Riolueang S, Glomglao W, et al. Diagnostic applications of newborn screening for alpha-thalassaemias, haemoglobins E and H disorders using isoelectric focusing on dry blood spots. Annals of clinical biochemistry 2013.

10. Karakochuk CD, Whitfield KC, Barr SI, et al. Genetic hemoglobin disorders rather than iron deficiency are a major predictor of hemoglobin concentration in women of reproductive age in rural prey veng, Cambodia. Journal of Nutrition 2015; 145(1): 134–42.

11. Karnpean R, Pansuwan A, Fucharoen G, Fucharoen S. Evaluation of the URIT-2900 Automated Hematology Analyzer for screening of thalassemia and hemoglobinopathies in Southeast Asian populations. Clin Biochem 2011; 44(10-11): 889–93.

12. Koh DXR, Raja Sabudin RZA, Mohd Yusoff M, et al. Molecular Characterisation of alpha- and beta-Thalassaemia among Indigenous Senoi Orang Asli Communities in Peninsular Malaysia. Annals of human genetics 2017.

13. LemmensZygulska M, Eigel A, Helbig B, Sanguansermsri T, Horst J, Flatz G. Prevalence of alpha-thalassemias in northern Thailand. Hum Genet 1996; 98(3): 345–7.

14. Limsakulsiriratt P, Oncoung W. High Risk Couples for Hb Bart’s Hydrops Fetalis in Public Health Region 8 and 9 During 2004-2006. Chonburi Hospital Journal 2007; 32(1): 9–14.

15. Munkongdee T, Pichanun D, Butthep P, et al. Quantitative analysis of Hb Bart’s in cord blood by capillary electrophoresis system. Ann Hematol 2011; 90(7): 741–6.

16. Munkongdee T, Tanakulmas J, Butthep P, et al. Molecular Epidemiology of Hemoglobinopathies in Cambodia. Hemoglobin 2016; 40(3): 163–7.

17. Nguyen HV, Sanchaisuriya K, Nguyen D, et al. Thalassemia and Hemoglobinopathies in Thua Thien Hue Province, Central Vietnam. Hemoglobin 2013; 37(4): 333–42.

18. Nguyen NT, Sanchaisuriya K, Sanchaisuriya P, et al. Thalassemia and hemoglobinopathies in an ethnic minority group in Central Vietnam: implications to health burden and relationship between two ethnic minority groups. Journal of Community Genetics 2017: 1– 8.

19. Nguyen VH, Sanchaisuriya K, Wongprachum K, et al. Hemoglobin Constant Spring is markedly high in women of an ethnic minority group in Vietnam: a community-based survey and hematologic features. Blood cells, molecules & diseases 2014; 52(4): 161–5.

20. Nillakupt K, Nathalang O, Arnutti P, Jindadamrongwech SB, T., Panichkul S, Areekul W. Prevalence and hematological parameters of thalassemia in Tha Kradarn subdistrict Chachoengsao Province, Thailand. J Med Assoc Thai 2012; 95(Suppl 5): S124–S32.

21. O’Riordan S, Hien TT, Miles K, et al. Large scale screening for haemoglobin disorders in southern Vietnam: implications for avoidance and management. British journal of haematology 2010; 150(3): 359–64.

22. Panomai N, Sanchaisuriya K, Yamsri S, et al. Thalassemia and iron deficiency in a group of northeast Thai school children: relationship to the occurrence of anemia. Eur J Pediatr 2010; 169(11): 1317–22.

23. Panyasai S, Cheechang S. The efficiency of screening for carriers of severe thalassemia in three community hospitals in Nakhon Si Thammarat province, Thailand. Songkla Med J 2009; 27(1): 61–72.

24. Pharephan S, Sirivatanapa P, Makonkawkeyoon S, Tuntiwechapikul W, Makonkawkeyoon L. Prevalence of α-thalassaemia genotypes in pregnant women in northern Thailand. Indian Journal of Medical Research 2016; 143(MARCH): 315–22.

25. Phollarp P, Tritipsombut J, Worasan C, et al. Thalassemia and iron deficiency among pregnant women attending antenatal care service at Khao Wong Hospital, Kalasin province. Journal of Medical Technology and Physical Therapy 2010; 22(3): 262–70.

26. Pichanun D, Munkongdee T, Klamchuen S, et al. Molecular Screening of the Hbs Constant Spring (codon 142, TAA > CAA, alpha 2) and Pakse (codon 142, TAA > TAT, alpha 2) Mutations in Thailand. Hemoglobin 2010; 34(6): 582–6.

27. Rahimah AN, Nisha S, Safiah B, et al. Distribution of alpha thalassaemia in 16 year old Malaysian Students in Penang, Melaka and Sabah. The Medical journal of Malaysia 2012; 67(6): 565–70.

28. Sanchaisuriya K, Fucharoen S, Ratanasiri T, et al. Thalassemia and hemoglobinopathies rather than iron deficiency are major causes of pregnancy-related anemia in northeast Thailand. Blood Cell Mol Dis 2006; 37(1): 8–11.

29. Savongsy O, Fucharoen S, Fucharoen G, Sanchaisuriya K, Sae-ung N. Thalassemia and hemoglobinopathies in pregnant Lao women: carrier screening, prevalence and molecular basis. Ann Hematol 2008; 87(8): 647–54.

30. Sengchanh S, Sanguansermsri T, Horst D, Horst J, Flatz G. High frequency of alpha-thalassemia in the So ethnic group of south Laos. Acta Haematol-Basel 2005; 114(3): 164– 6.

31. Setianingsih I, Harahap A, Nainggolan IM. Alpha thalassaemia in Indonesia: phenotypes and molecular defects. Advances in experimental medicine and biology 2003; 531: 47–56.

32. Sornkayasit K, al. e. Incidence of Hb Constant Spring and Hb Pakse in Khon Kaen: Using capillary electrophoresis and DNA analysis (Poster abstract). The 18th National Thalassemia Academic Symposium. Nonthaburi; 2012. p. 90.

33. Srivorakun H, Fucharoen G, Changtrakul Y, Komwilaisak P, Fucharoen S. Thalassemia and hemoglobinopathies in Southeast Asian newborns: diagnostic assessment using capillary electrophoresis system. Clin Biochem 2011; 44(5-6): 406–11.

34. Sutjasung P, Fucharoen G, Fucharoen S, Chattumaruk P, Changtrakun D, Sanchaisuriya K. Effectiveness of thalassemia screening with the use of internal quality control blood samples at Kasetsomboon Hospital, Chaiyaphoom province. Journal of Medical Technology and Physical Therapy 2011; 23(1): 34–45.

35. Suwannakhon N, Seeratanachot T, Mahingsa K, Namwong P, T. S. Prevalence of Alpha-thalassemia Trait in the Volunteered Personals of University of Phayao. J Hematol Transfus Med 2014; 24: 129–36.

36. Tan JA, Tay JS, Soemantri A, et al. Deletional types of alpha-thalassaemia in central Java. Hum Hered 1992; 42(5): 289–92.

37. Tan Jama, Lee PC, Wee YC, et al. High prevalence of alpha- and beta-thalassemia in the kadazandusuns in east Malaysia: Challenges in providing effective health care for an indigenous group. Journal of Biomedicine and Biotechnology 2010; 2010.

38. Tangvarasittichai O, Jeenapongsa R, Sitthiworanan C, Sanguansermsri T. Laboratory investigations of Hb Constant Spring. Clin Lab Haematol 2005; 27(1): 47–9.

39. Tangvarasittichai O, Poonanan N, Tangvarasittichai S. Using Red Cell Indices and Reticulocyte Parameters for Carrier Screening of Various Thalassemia Syndromes. Indian journal of clinical biochemistry: IJCB 2017; 32(1): 61–7.

40. Tanphaichitr VS, Pung-amritt P, Puchaiwatananon O, et al. Studies on hemoglobin Bart’s and deletion of alpha-globin genes from cord blood in Thailand (poster abstract). The International Conference on Thalassemia. Bangkok; 1985. p. P05.

41. Than AM, Harano T, Harano K, Myint AA, Ogino T, Okada S. High incidence of alpha-thalassemia, hemoglobin E, and glucose-6-phosphate dehydrogenase deficiency in populations of malaria-endemic southern Shan State, Myanmar. Int J Hematol 2005; 82(2): 119–23.

42. Tongon R, Yunu R, Sanchaisuriya K, et al. Thalassemia and hemoglobinopathies in pregnant women attended antenatal care service at Yala Hospital. J Med Tech Phy Ther 2014; 26(1): 32–9.

43. Tritipsombut J, Sanchaisuriya K, Fucharoen S, et al. Hemoglobin Profiles and Hematologic Features of Thalassemic Newborns Application to Screening of alpha-Thalassemia 1 and Hemoglobin E. Arch Pathol Lab Med 2008; 132(11): 1739–45.

44. Tritipsombut J, Sanchaisuriya K, Phollarp P, et al. Micromapping of Thalassemia and Hemoglobinopathies in Diferent Regions of Northeast Thailand and Vientaine, Laos People’s Democratic Republic. Hemoglobin 2012; 36(1): 47–56.

45. Uaprasert N, Settapiboon R, Amornsiriwat S, et al. Diagnostic utility of isoelectric focusing and high performance liquid chromatography in neonatal cord blood screening for thalassemia and non-sickling hemoglobinopathies. Clinica chimica acta; international journal of clinical chemistry 2014; 427: 23–6.

46. Yap ZM, Sun KM, Teo CRL, Tan ASC, Chong SS. Evidence of differential selection for the -alpha(3.7) and -alpha(4.2) single-alpha-globin gene deletions within the same population. Eur J Haematol 2013; 90(3): 210–3.

47. Yin SKK, Chong QT, Mei LA, et al. A molecular epidemiologic study of thalassemia using newborns’ cord blood in a multiracial Asian population in Singapore - Results and recommendations for a population screening program. J Pediat Hematol Onc 2004; 26(12): 817–9.

